# Dynamics of dominance: maneuvers, contests, and assessment in the posture-scale movements of interacting zebrafish

**DOI:** 10.1101/2023.11.21.567896

**Authors:** Liam O’Shaughnessy, Tatsuo Izawa, Ichiro Masai, Joshua W. Shaevitz, Greg J. Stephens

**Author notes:** JWS and GJS contributed equally to this work.

## Abstract

While two-body fighting behavior occurs throughout the animal kingdom to settle dominance disputes, important questions such as how the dynamics ultimately lead to a winner and loser are unresolved. Here we examine fighting behavior at high-resolution in male zebrafish. We combine multiple cameras, a large volume containing a transparent interior cage to avoid reflection artifacts, with computer vision to track multiple body points across multiple organisms while maintaining individual identity in 3D. In the body point trajectories we find a spectrum of timescales which we use to build informative joint coordinates consisting of relative orientation and distance. We use the distribution of these coordinates to automatically identify fight epochs, and we demonstrate the post-fight emergence of an abrupt asymmetry in relative orientations-a clear and quantitative signal of hierarchy formation. We identify short-time, multi-animal behaviors as clustered transitions between joint configurations, and show that fight epochs are spanned by a subset of these clusters, which we denote as maneuvers. The resulting space of maneuvers is rich but interpretable, including motifs such as “attacks” and “circling”. In the longer-time dynamics of maneuver frequencies we find differential and changing strategies, including that the eventual loser attacks more often towards the end of the contest. Our results suggest a reevaluation of relevant assessment models in zebrafish, while our approach is generally applicable to other animal systems.

## INTRODUCTION

Social interactions drive some of the most intriguing and complex aspects of animal behavior, from swarming and schooling in large groups [1] to courtship and fighting between pairs [2–4]. In our own species, the effective modelling of partners in teaching and conversation [5–7], as well as non-verbal improvisational games (see e.g. [8]), is a fundamental aspect of the human social experience. Abnormalities in these interactions are connected to dysfunction such as in autism (see e.g. [9]).

Qualitatively, the complexity of social behavior varies with the number of interacting animals. In the collective behavior of flocks, swarms and crowds, relatively simple interactions across a large number of individuals can result in large-scale order [10], but this often comes with a reduced individual repertoire. In contrast, the behavior of animals in pairs and small groups is both less explored and challenging in its dynamical irregularities, lacking an obvious canonical structure or order parameter. While there has been remarkable recent effort in posture-scale, quantitative approaches to understanding naturalistic behavior in single animals [11–15], settings with multiple organisms are substantially fewer (though see e.g. [16]).

Among the many intriguing facets of social behavior, those associated with the broad concept of dominance [17] are universally expressed in gregarious species. Dominance interactions produce and structure group hierarchies [18] and the position of an organism within a hierarchy has far-ranging consequences, from directly impacting the chance for reproduction, to health and disease [19, 20], to molecular and circuit-scale changes within the organism [21]. But dominance is an emergent relationship; one which animals must determine through their interactions. In fact, throughout the animal kingdom, from fruit flies [2], to shrimp [3], to elephant seals [4], dominance is established through a structured series of close and intense interactions, the collection of which is known as a “fight” or “contest” [22].

Animal contests are hypothesized to follow a general stereotyped structure [22, 23]. In this qualitative picture, contests begin with visual and/or vocal displays [24, 25], and then progress to physical interactions, which can fluctuate in intensity, until a dominance relationship is established [26]. Despite the appearance of a similar contest structure across species, *quantitatively* little is known about the organization of fights, the short-time behaviors that compose fights, and how contestants use these motifs to establish a dominance ordering over longer timescales. From high-resolution recordings how do we identify a contest? What, if any, are the differences between the beginning and the end of the contest? How do we identify dominance and can we find differences between the animals *before* the end of the contest? We address these fundamental questions by leveraging highresolution imaging and analysis of the dynamics of dominance contests in pairs of adult male zebrafish, *Danio rerio*.

Previous work has shown that pairs of male zebrafish engage in dominance contests when isolated in a novel environment [27–30]. In all of these efforts, however, the interactions were either scored by eye [28], or human-specified behaviors were identified using supervised machine learning [29]. Furthermore, current understanding of zebrafish contests is limited to broad differences between in-contest and post-contest behavior. But fight dynamics are fast and complex, and thus such approaches may capture only a subset of the full richness of the interactions, including how the interactions develop over the course of a contest. Here we combine quantitative analysis with 3D tracking of multiple body points across multiple organisms, obtained from a custom tracking apparatus consisting of multiple cameras, a large imaging volume, and a transparent interior cage to avoid reflection artifacts, to provide a high-resolution view of contest dynamics.

## RESULTS

### Imaging & Tracking

When animals interact they generally control both their egocentric pose as well as their location and orientation relative to others. Here we capture both the pose of individuals and their configurations in the environment by measuring the 3D coordinates of multiple bodypoints on multiple animals. In Fig. 1 we outline our image processing pipeline in which we track three body points on each of a pair of adult male zebrafish in three spatial dimensions, all while maintaining organism identity and without physical tags. In brief, we image fish behavior using three simultaneously-triggered cameras situated along orthogonal views of a 30 × 30 × 30 cm^3^ volume (∼ 10 body lengths per dimension) defined by a transparent interior cage constructed to avoid reflection artifacts, SI Fig. S1. In each view, we identify three body points, approximately corresponding to the tip of the head, the base of the pectoral fins, and the tail of each zebrafish. We use a custom implementation of the SLEAP algorithm [31] to automatically recognize these “head”, “pec” and “tail” body points in each view throughout the recording, Fig. 1(A). From the set of 2D views, we compute the 3D position of each body point in lab coordinates using a calibrated camera model, SI Fig. S2. To maintain organism identity, we apply a separate machine vision solution, idtracker.ai [32] to the top-down camera view and register idtracker.ai identity labels with 3D SLEAP skeletons using the Hungarian sorting algorithm [33]. The output of our tracking pipeline is thus the 3D coordinates for the three body points of each organism. Further tracking details are provided in Methods, and see SI Mov. S1 for a visualization of the tracking pipeline output. Our particular body point choices enable the identification of animal location, which we define as the “pec” point, animal orientation, which we define as the direction of the unit vector from “pec” to “head” points, as well as the dynamics of the tail. In the social conditions studied here, the closely interacting fish move throughout all three dimensions of the imaging volume, Fig 1(B), are often separated by less than a few body lengths, and are strongly correlated in their orientations, Fig. 1(C).

**FIG. 1.**
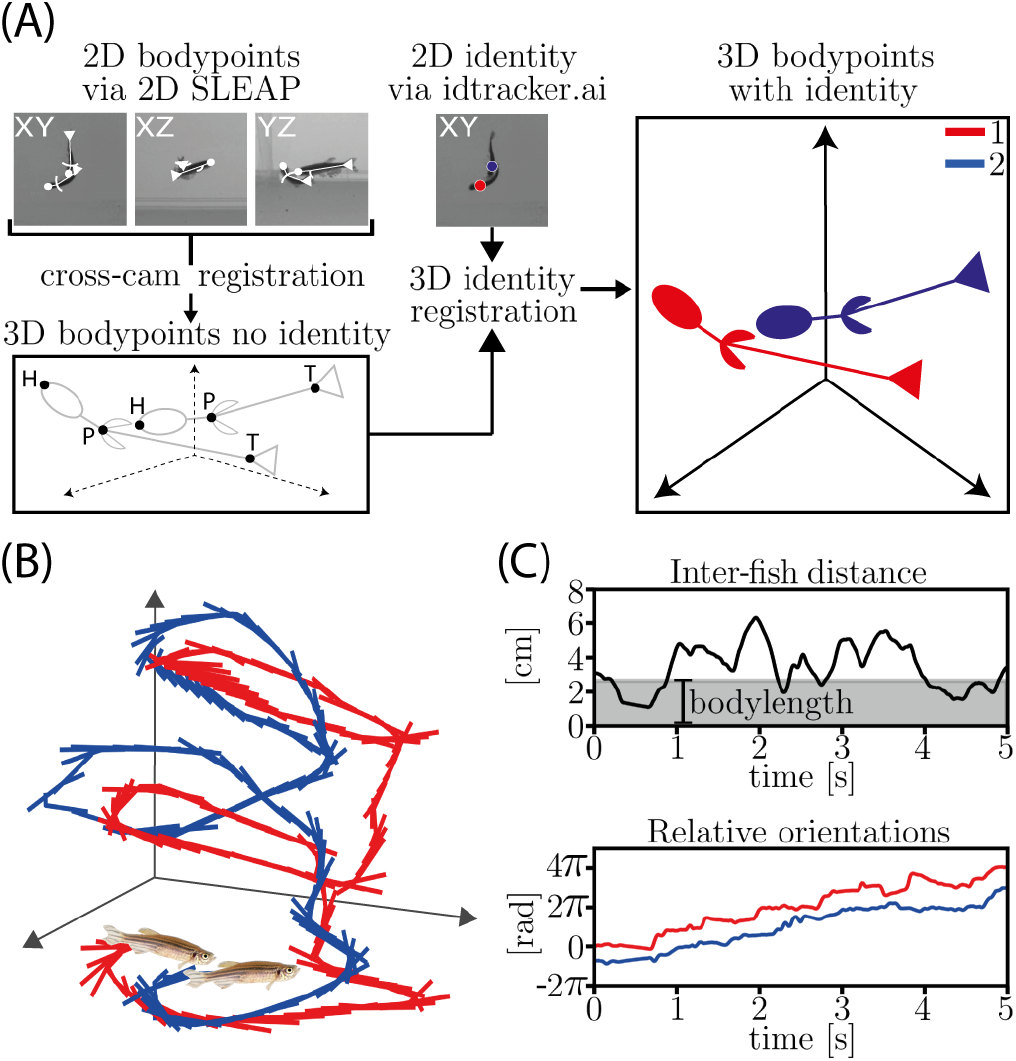
Markerless keypoint tracking of strongly interacting zebrafish in 3D. (A) Schematic of the tracking pipeline. We record images from three orthogonal views of a 40 cm × 40 cm × 44 cm tank (SI Fig. S1). Using SLEAP [31], we localize three keypoints from each animal within the 2D images: the tip of the head, the base of the pectoral fins, and the base of the tail. We find the 3D position of each keypoint from the set of 2D views using a calibrated camera model, SI Fig. S2. Separately, we maintain animal identity over time using idtracker.ai [32] on the XY camera view. For the final tracking output, we combine fish identity (labelled in red and blue) with the 3D pose. (B) Example reconstructed 5 s trajectory of the 3D position of the two fish. For every fifth time point in the trajectory we draw a skeleton connecting the three keypoints. (C) State variables used in subsequent analysis for the segment shown in panel B. (C-top) The interfish distance, defined as the 3D euclidean distance between pectoral keypoints, with the average body length (gray) for scale. (C-bottom) Planar relative orientations of the two fish defined as the direction of each pec-head vector projected onto the XY-plane (see also Fig. 2 and SI Fig. S4). Strong interactions are apparent in their close distance and aligned but changing orientation. See SI Mov. S1 for the visualization of an entire recording.

### Interpretable Joint Coordinates

Our tracking provides an 18-dimensional instantaneous representation of two-fish configurations. We expect that the behavior is approximately invariant to location and planar direction within the imaging volume. Thus, we eliminate four of the dimensions corresponding to translations of the joint-configuration centroid location and the joint overall orientation within the xy-plane. Within the remaining dimensions, we find that further simplifications are naturally apparent from principal components analysis, Fig. 2(A). Ordered by their fraction of captured variance, the first two modes (orange) describe configurations differing in xyand zseparations between the two fish, respectively. Both separations can be approximately as large as the linear dimension of the interior cage volume. The next six modes (green) characterize relative changes between the pitch and yaw of each organism. In our conditions, the pitch angle is always relatively small, but yaw can vary fully around the circle. Thus four linear dimensions are needed to account for this nonlinear yaw variation of the two organisms. The last six modes account for changes in the posture (egocentric pose) of each animal.

**FIG. 2.**
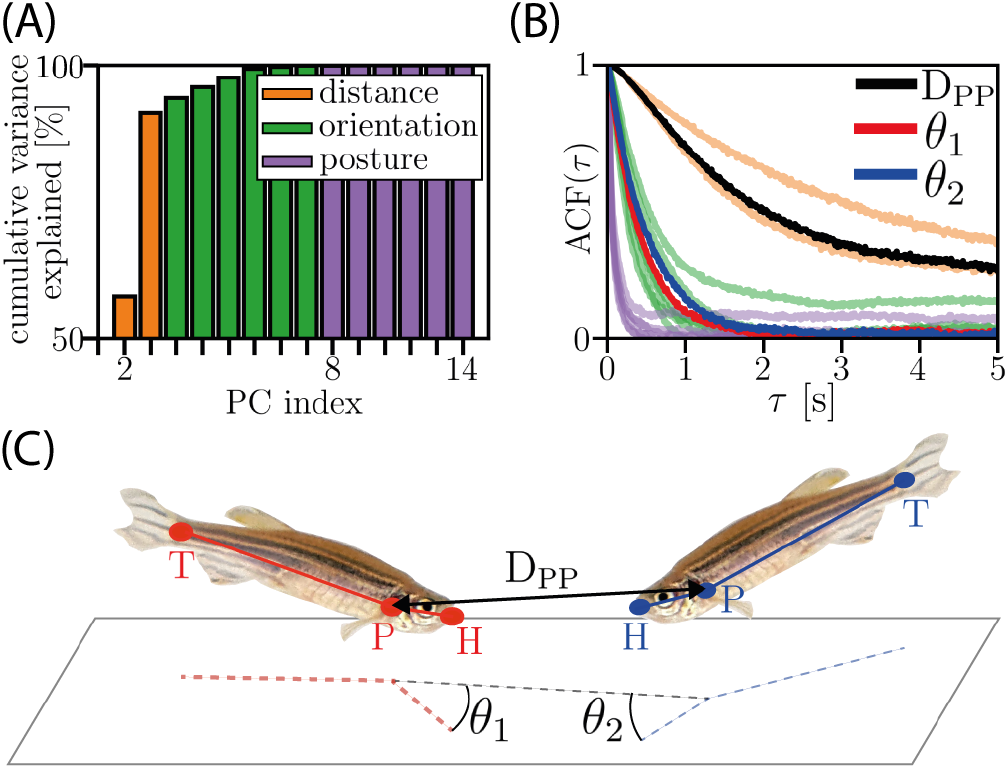
Two-body state variables. (A) Principal components analysis of keypoint positions in the co-rotating frame reveals 3 sets of modes: (orange) relative distance, (green) relative orientation, and (purple) posture deformation. The two distance modes correspond to separation along the horizontal and vertical directions, while the six orientation modes correspond to four modes for the linear basis of the two relative orientations *θ*_1_, *θ*_2_ and two small amplitude pitch angles. (B) A separation of timescales is apparent in the temporal autocorrelation of the PCA modes: relative distances change slowly, relative orientations are faster, and posture deformations exhibit the fastest dynamics. (C) We collapse the 14dimensional set of co-rotating coordinates to 3 interpretable variables that cover the distance and orientation timescales: *D*_*PP*_ (the distance between the pectoral keypoints, black), *θ*_1_ (the relative orientation of fish 1, red), and *θ*_2_ (the relative orientation of fish 2, blue), see also SI Fig. S4. We leave the fast posture modes as well as the vertical separation and pitch angle for future analysis. We also find that the dominance contests are primarily planer with the joint centroid drifting upwards, SI Fig. S5, and that the pitch angles are strongly peaked around zero, SI Fig. S6.

We find that the joint configuration modes vary with distinct timescales, Fig. 2(B). We normalize each mode to unit variance and compute the auto-correlation function *ACF* (*τ*) = ⟨ (*x*(*t* + *τ*) − ⟨*x*(*t*) ⟩)(*x*(*t*) −⟨*x*(*t*) ⟩) . Variations in separation are slowest, reorientation dynamics are faster, while changes in posture are the fastest of all. We use this timescale ordering to motivate a simplified and interpretable coordinate system in which we capture the separation modes through a single pec-pec distance *D*_*P P*_ and the orientation modes in the relative angles *θ*_1_ and *θ*_2_, Fig. 2(C). We leave the fast posture modes as well as the explicit z-separation and pitch angle for future analysis. We show later that dominance contests are primarily planer with the joint centroid drifting upwards, SI Fig. S5, and that the pitch angles are strongly peaked around zero, SI Fig. S6. We additionally compute the XY rotational velocity 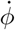 of the two-fish system, where *ϕ*(*t*) is the angle between *X*_*Lab*_ and *X*^*′*^, SI Fig. S4. We take the instantaneous state space spanned by {*D*_*P P*_, *θ*_1_, *θ*_2_} as the basis for further analysis of the contest dynamics.

### Fight bout Detection and Winner/Loser Determination

In both zebrafish and other organisms, contest epochs have been typically identified by eye [2, 3, 30], but we seek to systematically discover these epochs using the configuration coordinates {*D*_*P P*_, *θ*_1_, *θ*_2_}. In Fig. 3(A) we present the dynamics of a typical experiment, viewed though the probability of each configuration coordinate, computed in running 1-minute windows, a duration long compared to the correlation time of each variable, but short enough to still reveal important dynamics. Within our recordings there may be none, one, or multiple epochs of active fighting (hereafter called fight bouts). Vertical dotted lines denote fight bout boundaries computed using the clustering described below and fight bout epochs are clearly visible. For a visualization of the dynamics before, during and after these fight bouts, see SI Mov. S2-S6.

**FIG. 3.**
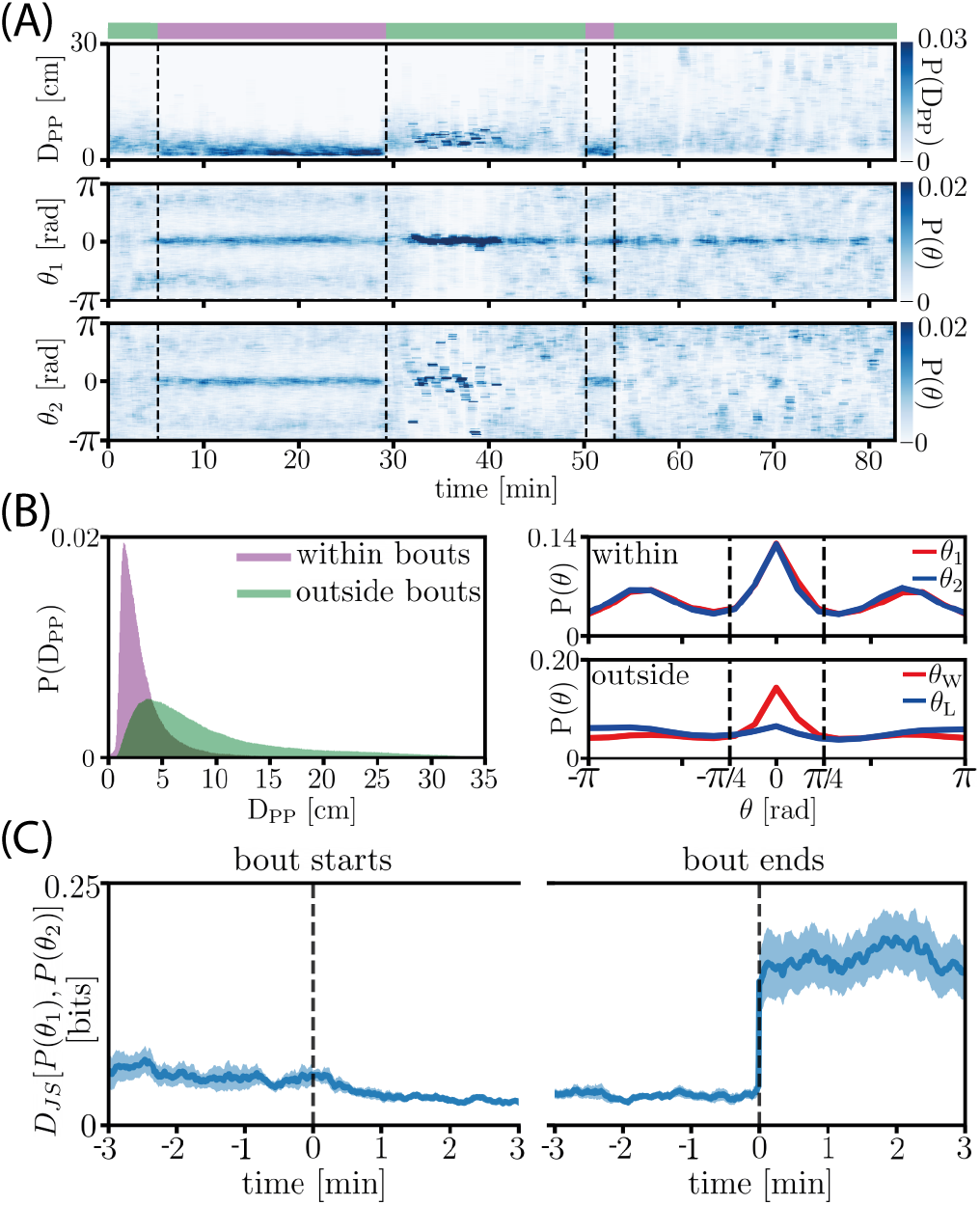
Identification of fight bout epochs and winner/loser assignment. (A) We show the distributions *P* (*D*_*PP*_), *P* (*θ*_1_), and *P* (*θ*_2_) for a single recording computed in sliding windows (1 min length, 59 sec overlap). At the coarsest scales, we observe two regimes, corresponding to fighting and non-fighting (SI Fig. S7). Fighting occurs when the animals are close (small *D*_*PP*_) and in orientations conducive to chasing and attacking (peaks in *θ*_1_, *θ*_2_ at 0 and *±*0.8*π* rad). (B-top) The distribution *P* (*D*_*PP*_) during and outside of fight bouts. (B-middle) The distributions *P* (*θ*_1_) and *P* (*θ*_2_) during fight bouts, illustrating a symmetry between contestants. (Bbottom) The distributions for the winner and loser, *P* (*θ*_*W*_) and *P* (*θ*_*L*_), are asymmetric after the fight ends. We assign winner and loser labels to each contestant through the post-fight angular distributions in which losers have lost their strong peak at *θ* = 0 (see also SI Fig. S9). (C) The JensenShannon divergence *D*_*JS*_ [*P* (*θ*_1_), *P* [*θ*_2_]) aligned to fight bout starts (left) and to fight bout ends (right). We plot the mean (dark line) and standard deviation (shaded region) for fight bouts longer than 7 minutes. The symmetry in orientation (small *D*_*JS*_) disappears abruptly when fight bouts end. See SI Mov. S2-S6 for visualizations of the dynamics before, during and after fight bouts.

Quantifying the intuition from this exemplar recording, we develop a fight bout “detector” by clustering the windowed, 3D joint configuration distributions *P* (*D*_*P P*_, *θ*_1_, *θ*_2_). We compute this probability after passing the *θ* variables through a symmetric function to avoid making a choice between the labelling of *θ*_1_ and *θ*_2_. We organized the resulting collection of window distributions using hierarchical clustering with the square root of the Jensen-Shannon divergence *D*_*JS*_ as a metric, and identified a clear cluster corresponding to fight bouts, SI Fig. S7. In our *N* = 22 recordings we identified *N*_bouts_ = 45 fight bouts with mean duration of 11 min (we show the distribution of detected fight durations in SI Fig. S8). Full clustering details are provided in Methods. In Fig. 3(B) we apply our fight bout detector across the entire set of recordings to show that fight bouts are stereotyped configurations characterized by small separations, *D*_*P P*_ ∼ bodylength, and orientation angles strongly peaked towards the other fish, *θ* ≈ 0.

The example in Fig. 3(A) also illustrates an important change in the joint configurations which occurs after fight bouts: one animal maintains a relative orientation distribution with a strong peak at *θ* ≈ 0, while this peak is greatly diminished in the other animal. Thus a symmetry between the two animals has been broken. We use this broken symmetry to quantitatively identify a “winner” and “loser” in each recording (which again, may contain multiple fight bouts). We note that the behavior of the contestant with a strong, post-contest peak at *θ* ≈ 0 is qualitatively consistent with previous observations of continuing aggression [30, 34]. We thus make a winner/loser determination by computing the summed probability in the 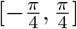 regions of the *θ* distributions after fight bouts, Fig. 3(B-bottom). Across our full dataset, 20 out of 22 recordings revealed a clear winner, and the two experiments which did not reveal a clear winner were withheld from subsequent analysis. An additional 2 other experiments showed a clear winner from the beginning of the recording, meaning we could not study the emergence of winner and loser, and so we also withheld these two experiments from subsequent analysis, leaving N=18 experiments.

The symmetry in orientation distributions during fight bout epochs appears and disappears rapidly. We compute *D*_*JS*_ between *P* (*θ*_1_) and *P* (*θ*_2_) (both probabilities again sampled in 1-minute windows) and show the average across all fight bouts, aligned with the beginning and end of a bout, Fig. 3(C). In both cases, we examined 3 minutes before and after bout-start and bout-end, using all fight bout data longer than 7 minutes. For bout endings (right panel) we find a sudden transition from a symmetric epoch with small *D*_*JS*_ between distributions, to an asymmetric epoch with larger divergence. For bout beginnings this transition is more gradual (left panel).

### Stereotypy in Orientation Dynamics

The clear interpretation of the relative orientations {*θ*_1_, *θ*_2_} suggests that these variables also offer a window into the short-time dynamics that make up a bout. In Fig. 4(A) we show a schematic guide to the joint configurations corresponding to the values of each relative orientation. We then display the joint configuration probability *P* (*θ*_*W*_, *θ*_*L*_) computed within fight bouts Fig. 4(B, left) and for non-fight epochs, Fig. 4(B, right). Both distributions have informative structure. Outside of fight bouts, where our sampling is dominated by the post-fight epoch, we find signatures of dominance as the winner fish orients towards the loser but the loser orients away from the winner. These correspond to the peaks {*θ*_*W*_ ≈ 0, *θ*_*L*_ ≈ *π*}. There are two approximately symmetric peaks corresponding to whether the winner points to the right or left side of the loser fish, respectively. Also notable is a peak in the center ({*θ*_*W*_ ≈ 0, *θ*_*L*_ ≈ 0} reflecting a “face-off” configuration in which both fish are oriented directly towards each other. Within fight bouts the structure of *P* (*θ*_*W*_, *θ*_*L*_) is quite different; we see prominent peaks corresponding to aggressive orientations from both fish, where a fish is directly facing its opponent, as well as a more continuous density connecting these peaks, perhaps reflective of transitions between aggressive states. The structure apparent in *P* (*θ*_*W*_, *θ*_*L*_) during fight bouts suggests the existence of short-time, stereotyped behaviors in the contest dynamics.

**FIG. 4.**
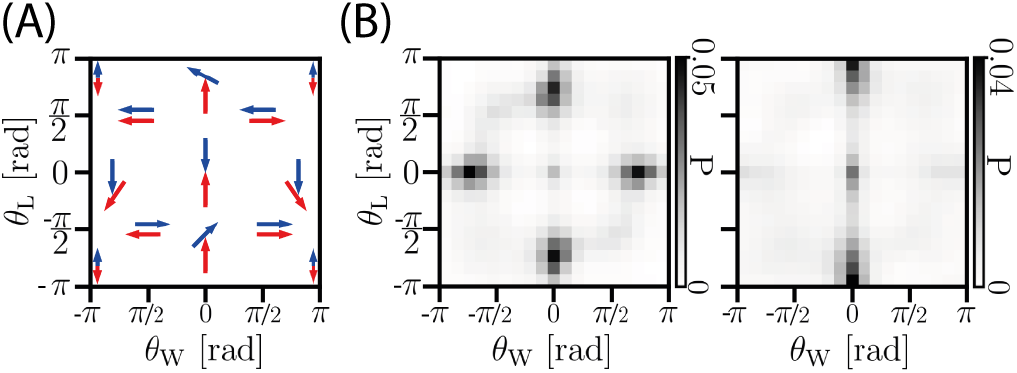
The joint distribution *P* (*θ*_1_, *θ*_2_) reveals stereotyped orientation dynamics. (A) Schematic illustrating possible relative orientations in the {*θ*_*W*_, *θ*_*L*_} space. (B) *P* (*θ*_*W*_, *θ*_*L*_) sampled from different epochs in our recordings. (B-left) Fight bouts. Peaks in the distribution are indicative of chasing and attacking by both winner and loser fish. (Bright) Non-fight epochs. Peaks show the winner fish oriented towards the loser, with the loser facing away from the winner.

### Identifying Fight Maneuvers

To quantify the stereotyped structure of contest dynamics, we constructed a transition matrix among discretized microstates of (*D*_*P P*_, *θ*_*W*_, *θ*_*L*_) configurations. We chose *N* = 20 evenly-spaced partitions for each variable, resulting in 8,000 microstates. We then built a transition matrix *T*_*ij*_ by counting transitions between microstates every *τ* = 10 frames (see Fig. 5(A-left) for a schematic illustration). We discuss the choice of *τ* in SI Fig. S10(A), and later in this section after we introduce our clustering process.

**FIG. 5.**
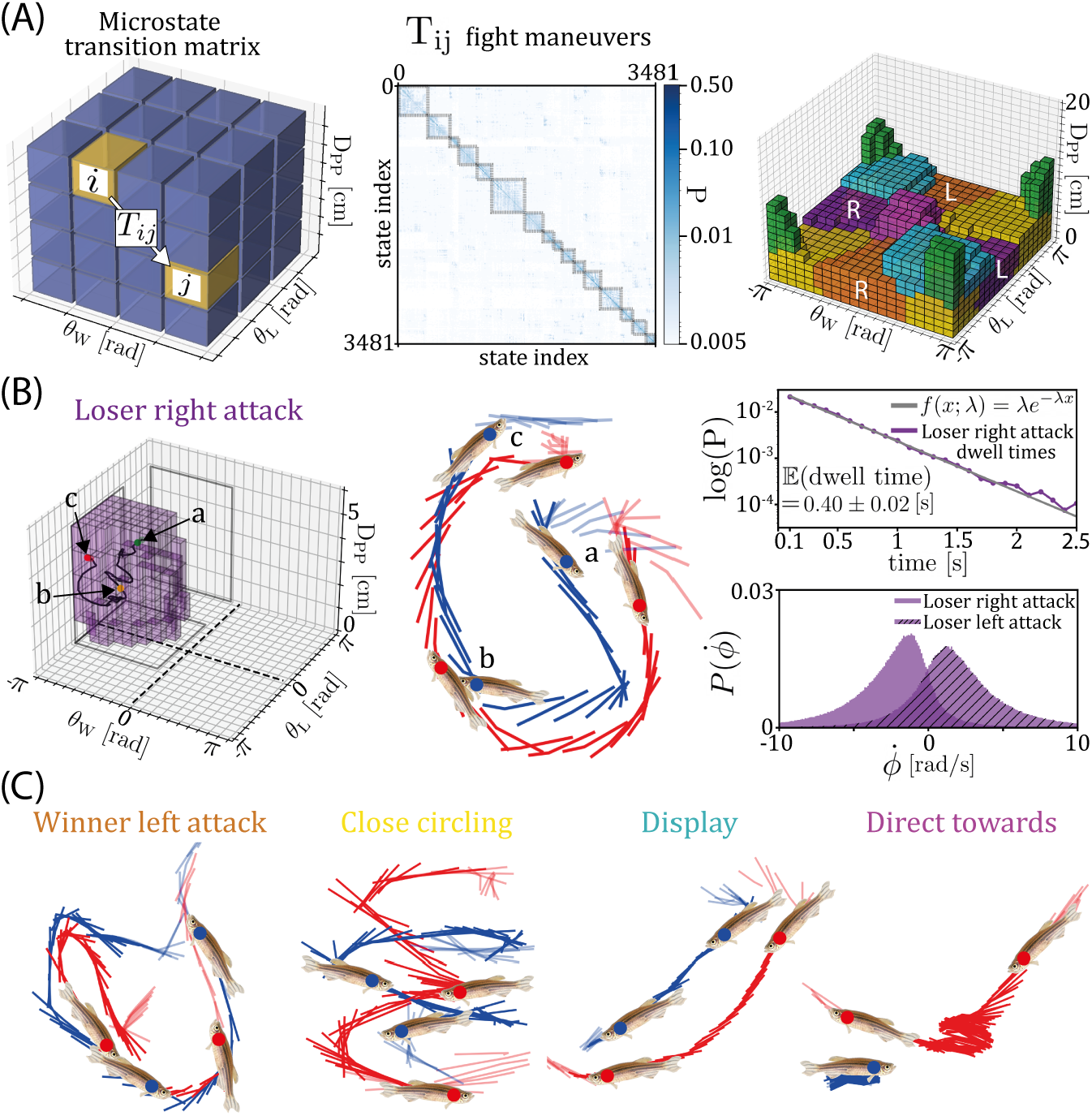
We identify fight maneuvers by clustering dynamical transitions between two-body configurations. (A) We use 20 equally-spaced partitions for each joint variable (*D*_*PP*_, *θ*_*W*_, *θ*_*L*_) and compute the transition matrix *T*_*ij*_ of all pairwise transition probabilities between the 8000 joint microstates. *T*_*ij*_ is a weighted directed graph and we use Infomap network community detection to identify *N* = 56 network communities, interpretable as dynamically clustered regions of the two-body space. From these we select a smaller (*N* = 16) set which span fight bouts (SI Fig. S10 (B)), and we further combine these through symmetry to arrive at *N* = 10 primary clusters, which we label as fight maneuvers (SI Fig. S11). (A-left) Schematic illustrating the computation of the transition matrix *T*_*ij*_ . (A-middle) We visualize a subset of the transition matrix *T*_*ij*_ containing only microstates which are members of the 16 network communities which span fight bouts. Rows of *T*_*ij*_ are sorted so the microstates belonging to each community are contiguous, illustrating a roughly block-diagonal structure in *T*_*ij*_ uncovered by Infomap. (A-right) We show the microstates belonging to six of the identified fight maneuvers, with the other four maneuvers excluded for visual clarity. (B-left) An example maneuver in which the loser attacks the right-hand side of the winner. We show an example trajectory through the cluster (black line), and also a visualization (B-middle) of the associated 3D trajectory. The lighter trajectory lines represent short time segments before and after the maneuver activation. (B-right, top) Maneuvers are short-lived with an approximately exponential dwell-time distribution. (B-right, bottom) The distribution of the two-body rotational velocity 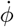 (see SI Fig. S4) reveals a clockwise bias for right attacks and a counterclockwise bias for left attacks. (C) Example 3D trajectories of four other fight maneuvers; label colors are consistent with (A-right).

The transition matrix encodes contest dynamics as a directed graph connecting microstates of (*D*_*P P*_, *θ*_*W*_, *θ*_*L*_) configurations. We coarse-grained this space by identifying groups of microstates which tend to transition between each other. We applied the Infomap network clustering algorithm [35], and obtained a set of *N* = 56 clusters. Each cluster is a collection of (*D*_*P P*_, *θ*_*W*_, *θ*_*L*_) microstates (see Fig. 5(A-right) and also SI Fig. S10(C)), and represents a coarse-grained macrostate of the two-fish system. Clusters are composed of contiguous microstates as the dynamics vary smoothly between different (*D*_*P P*_, *θ*_*W*_, *θ*_*L*_) configurations.

While approximately discrete behavioral macrostates were already apparent in Fig. 4, the number of clusters does depend on the transition time *τ* . Roughly when *τ* is small then every microstate is its own cluster as there is no time to transition away, and when *τ* is large then all microstates are connected into one cluster. We seek an intermediate *τ* which allows for an expressive number of clusters but where the clusters themselves are composed of multiple microstates. We note that the choice of a single transition time *τ* does not preclude multiple timescale dynamics; even an equilibrium Markov chain can exhibit myriad relaxation times.

To focus on the dynamics of fighting behavior we identify a subset of clusters which span fight bouts. We rankorder the *N* = 56 clusters by their occurrence frequency during fight bouts, and select the top *N* = 16 clusters which together cover 95% of frames during fight bouts (see SI Fig. S10(B)). In Fig. 5(A-middle) we show the transition matrix restricted to the 3481 microstates contained within these clusters. The rows and columns of the transition matrix are ordered by cluster membership, and exhibit a block diagonal structure. The *N* = 16 clusters capture interpretable course-grained dynamics which occur during fighting, and we denote these as “fight maneuvers”. We show the microstates composing these fight maneuvers in SI Fig. S10(C).

In Fig. 5(B) we examine the fight maneuver “Loser right attack”. In Fig. 5(B, left) we show the collection of microstates that make up the “Loser right attack” fight maneuver. All of these microstates occur at *D*_*P P*_ *<* 5 cm, meaning that this is a close-distance maneuver. The manuever microstates are centered around *θ*_*L*_ = 0 rad, meaning that the (eventual) loser is pointing towards the (eventual) winner, and the microstates have *θ*_*W*_ strictly negative and centered around 0.8 rad, which represents the winner pointing away from the loser. Shown in black is the trajectory in (*D*_*P P*_, *θ*_*W*_, *θ*_*L*_) corresponding to an exemplar activation of this fight maneuver. We plot the 3D coordinates of the fish during this same maneuver activation in Fig. 5(B, middle). The pair begin at a distance of several centimeters (point a). Then the loser closes the distance to the winner and makes a contact from behind (point b). Finally the pair move apart again (point c). In Fig. 5(B, right) we present additional properties of this maneuver. In the top panel we show the distribution of dwell times (purple), which are approximately exponentially distributed (fit, grey) in the range of 0 − 2.5 s, with a mean of 0.40 *±* 0.02 s, where the error is the standard deviation from bootstrapping across experiments. In the bottom panel we show the distribution of the XY rotational velocity 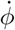 of the two-fish coordinate system (see SI Fig. S4), conditioned on activation of the “Loser right attack” maneuver, and conditioned on activation of the “Loser left attack” maneuver. During right attacks the fish pair spin clockwise, and during left attacks they spin anti-clockwise.

In Fig. 5(C) we plot 3D trajectories during an exemplar activation of four other fight maneuvers. We show in red a “Winner left attack”, where the winner attacks the left hand side of the loser. Next we show in yellow a maneuver we call “Close circling”. In this maneuver the fish are orientated in an anti-parallel configuration. 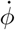 conditioned on this maneuver is large, representing spinning of the system. We use the term “close” as this maneuver happens at relatively small *D*_*P P*_ . Taken together, close distance and anti-parallel spinning lead to the name “Close circling”, although the activation of this maneuver does not require the pair to trace a full or multiple circles. In orange we present an example of the maneuver we label “Close parallel”. In this close distance maneuver the pair orients in parallel, and at close distance. Finally in purple we illustrate a maneuver we call “Direct towards”. In this close distance maneuver, the fish are orientated directly towards each other. Activation of this maneuver can have one or both of the fish moving slowly or stopped.

### Expression of Fight Maneuvers

For clarity we combined maneuvers which differ only in their left/right configuration, these are denoted by the same color in Fig. 5(A, right). After the left/right combination there are *N* = 10 maneuvers and we show their associated microstates in SI Fig. S11. In SI Fig. S12(A) we show an example ethogram (a time series of expressed behaviors) for the fight maneuvers over a 10 s period. This time period is an order of magnitude longer than the average dwell time of the maneuvers, but is short compared to the duration of fight bouts. We present the 6 most frequently expressed maneuvers in this time period, for which the complexity and speed of the behavioral dynamics is readily visible. For example, winner attack dwell times vary from ∼ 0.1 − 2.5 s (see Fig. 5(B, upper right)). Longer attack activations would be readily visible by eye, but we also see brief activations which would be difficult to identify without a quantitative approach.

We illustrate contest dynamics on a longer timescale in SI Fig. S12(B), where we show the progression of an exemplar experiment as seen through probability of the fight maneuvers over time. As fight bouts progress, there is a clear increase in the expression probabilities of winner and loser attacks. This is suggestive of an escalating aggression during the bouts [22, 36]. After both fight bouts, we find a significant difference in winner/loser attack probabilities. A drastic decrease in the loser attack probability illustrates the established dominance relationship between the pair. However, the period after the first and second fight bouts show differences. After the first fight bout, in the ∼ 20 min before the contestants fight again, we see periods of large “winner approach” and “direct towards” activations which we do not see after the second fight bout. After the second fight bout, where no further fighting is observed until the end of the recording, we see significantly more “direct away” activations than during the fight bouts or after the first fight bout. The expression of fight maneuvers viewed in this manner provides insights into the complexity of the dominance dynamics over (∼ hour) timescales, longer than those of both individual fight bouts (∼ 10 min) and fight maneuvers (∼ 1 s).

The fight maneuvers were chosen as a set of dynamically coherent clusters of (*D*_*P P*_, *θ*_*W*_, *θ*_*L*_) configurations which span fight bouts. In Fig. 6(A) we show the difference between fight bouts and non-fighting epochs as seen through the probability of these maneuvers. During fights, winner and loser attacks are approximately equally expressed. Outside fight bouts, attack probabilities are greatly decreased, with a significant difference between winner and loser. The probability of “close circling” is much higher during fights. The “winner approach” maneuver increases in probability outside of fights where there is a clear dominance relationship with the winner chasing the loser. As expected, the probability of non-fight-clusters, which we define as the compliment of the set of the *N* = 56 original clusters with the *N* = 16 fight maneuvers, is much higher outside of fights. The error bars are the standard deviations from bootstrapping across experiments.

**FIG. 6.**
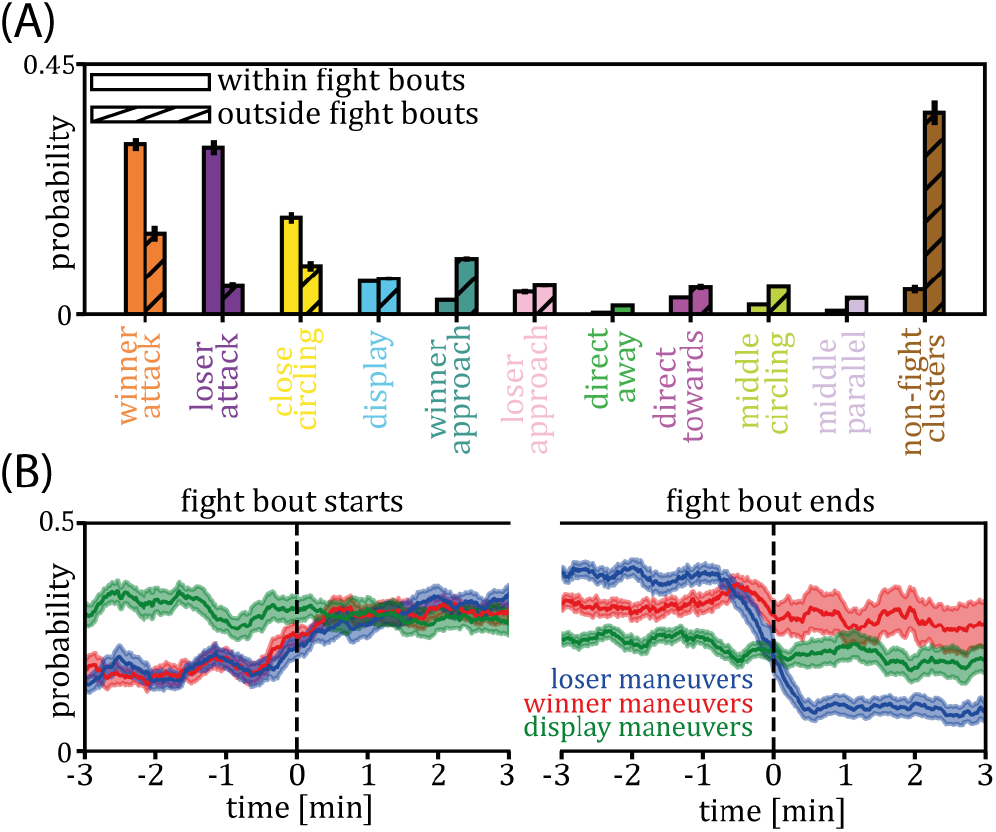
Maneuver probabilities across time suggest a loser component of the dominance decision. (A) Maneuver probabilities sampled from within (plain bars) and outside (hatched bars) fight bouts. (B) Changes in maneuver probabilities between the start and end of fight bouts. To focus on the asymmetry between contestants, we combine fight maneuvers into three categories: winner-advantageous, loser-advantageous and those which are symmetric between contestants. In detail, we combine “winner attacks” and “winner approaches” into “winner maneuvers”, and similarly for “loser maneuvers”. We combine “close circling”, “far circling”, “close parallel” and “far parallel” into “display maneuvers”. We then examine the windowed probabilities of these maneuvers aligned to the beginning and end of fight bouts. Probabilities are estimated in 30-sec running windows with 29-sec overlap. Plotted is the mean (dark line) and bootstrapped standard deviation (shaded region) from fight bouts longer than 7-minutes. (Left) Before the onset of fight bouts, “display maneuvers” are most probable, consistent with the concept of a display-like phase early in contests where aggression between contestants has yet to peak. (Right) After the end of the fight bouts, there is a clear asymmetry in “winner attacks” relative to “loser attacks”, highlighting the decisive outcome of the contest. Remarkably, in the minutes preceding the end of the fight bout there is a small but significant contestant asymmetry;”loser maneuvers” are more likely than “winner maneuvers”, suggesting a role for the loser in the dominance decision. This bias for loser advantageous maneuvers is also apparent when computed from individual bouts across the entire ensemble of contests, SI Fig. S13.

### Starting & Ending fight bouts

In Fig. 6(B) we examine the trends in the expression probabilities of the fight maneuvers at the start and end of fight bouts. To focus on the emergent asymmetry between winner and loser, we simplified our fight maneuver set into winner maneuvers, loser maneuvers, and display maneuvers. We combined “winner attack” with “winner approach” into “winner maneuvers”, and “loser attack” with “loser approach” into “loser maneuvers”. We also combined the symmetric states “close circling”, “far circling”, “close parallel” and “far parallel” into “display maneuvers”.

We plot the probabilities of these simplified maneuvers triggered on the start and end of fight bouts. Before fight bouts end, loser maneuvers are on average more probable than winner maneuvers. Probabilities are estimated in running 30-s windows with 29-s overlap, and errors are standard deviations from bootstrapping across experiments. This bias for loser advantageous maneuvers is also apparent when computed from individual bouts across the entire ensemble of contests, SI Fig. S13. Towards the end of the contest, the fish that ultimately loses is actually attacking more. This asymmetry abruptly switches at the end of the fight. Regardless of the particular assessment process, the contest is simply not a marquee boxing match where the loser gets beaten into submission. We return to this point in the discussion.

## DISCUSSION

Within the rich and complex space of social interactions, pairwise fighting behavior provides a tractable and relevant example. Contests are highly structured and relatively long-lasting, exhibit repeated behavioral patterns, and have a clear start and finish, as well as an emergent symmetry-breaking between the two contestants. Usefully, contestants are strongly engaged by the dynamical stimuli provided by the other animal, thus reducing the impact of the artificial imaging environment necessary for high-resolution tracking.

To measure such strongly interactive behavior, we described a large imaging arena and computer vision workflow to track multiple bodypoints of multiple adult zebrafish in 3D while maintaining individual identity. We then analyzed pair dominance contests within a set of long recordings sampled at high temporal resolution. We believe that the origin of the contest behavior in our experiments is to establish control of the imaging arena [37]. In the wild, territorial males are known to aggressively stop other males from accessing spawning sites [38, 39]. Zebrafish also live in groups for which contests may help establish a dominance hierarchy [40–42].

We followed three points in 3D along each fish (“head”, “pec” and “tail”) and used these points to construct the joint configuration variables (*D*_*P P*_, *θ*_*W*_, *θ*_*L*_). We used distributions of these variables to automatically identify fight bout epochs, and to demonstrate an emergent and abrupt asymmetry in the post-fight distribution of relative orientations. The identified fights were long lasting, with a minimum of a few minutes but a maximum of almost 30 minutes, SI Fig. S8. We note that our large imaging volume allows ample room to flee, thus reducing the contestants immediate physical danger, and perhaps prolonging the intense interaction.

The orientation asymmetry allows us to identify the ultimate “winner” and “loser” in each recording; “winner” denotes the dominant individual as determined by their strong propensity to orient towards the other fish throughout the post-contest epoch. This asymmetry was clear in the majority of recordings and was not present before the contest. But how do the contests determine this dominance ordering? To address this question, we used the tracked trajectories to identify a set of coarsegrained, joint dynamical states, which we denote as “maneuvers”. Maneuvers arise from clustering the transitions between *D*_*P P*_, *θ*_*W*_, *θ*_*L*_ configurations, resulting in dynamical motifs (∼ 1s) that segment the fight dynamics into a small number of interpretable two-fish behaviors. We used the expression of these maneuvers over time to unravel contest structure.

We found that a set of *N* = 10 maneuvers was sufficient to capture the behavioral complexity of the contests (SI Fig. S10 (B) & SI Fig. S11) and that many of these maneuvers are directly interpretable, including winner and loser attacks. The evolution of the occupancy of these joint maneuvers reveals that, on average and towards the end of the fight, the ultimate loser attacks more frequently than the ultimate winner, even as attacks are energetically costly and can result in injury. Interestingly, an analogous observation has been made in cichlid contests [43]. In cichlid contests where neither contestant had existing ownership of the arena, fish which were predicted to lose by expert observers, and did indeed eventually lose, still escalated aggression to damaging levels. The “desperado effect” [44] describes such dynamics, where prospective winners tend not to escalate due to the high costs involved and prospective losers continue to fight and even escalate because they lack alternative strategies.

More generally, game theoretic analyses of animal contests have focused on models of pairwise assessment [45], but these are difficult to align to real contests. Central to such models are the concepts of *resource holding potential* (RHP), the ultimate ability of an individual to win a contest [22, 46], and *contest cost* which is incurred by individuals for engaging in a fight [45]. While measurable correlates for these concepts are unknown, a commonly used proxy for RHP is contestant size, which has been shown to effect contest duration, see e.g. [37]. In the experiments presented here, contestant size was approximately equal, removing this source of variation across recordings. Contest cost may be reflected in the fight duration and the number of attacks, but this is a tangled combination of (at least) the costs of receiving physical blows and the energy expenditure of maneuvering.

An interesting consequence of assessment models is the existence and sequence of contest phases, longer-lasting epochs of differentiated patterns of maneuvers, see e.g. [3]. Using the fight bout detector we can reliably divide our data into fighting and non-fighting epochs (Fig. 3) and the recordings with multiple fighting epochs demonstrate that the contests do not proceed unidirectionally through these phases. Are there finer graduations to the fight bouts themselves? The windowed maneuver probabilities of Fig. 6(B) reveal that display-like maneuvers are most probable just before the onset of bouts while attack probabilities are symmetric at the start but asymmetric towards the end of a bout. In future work it will be interesting to explore whether this variation in maneuver probabilities is more discrete, as suggested by the phases of assessment models, or continuous, perhaps reflecting a non-stationary update of relative strength.

Our { *θ*_*W*_, *θ*_*L*_, *D*_*P P*_ } reduction of the three-point body tracking is powerful, but certainly a simplification from the full 14 degrees of freedom available in the three-point representation, as well as from a richer tracking of both posture and appearance. In future work, it will be informative to measure the full curvature of the caudal fin allowing for studies of the tail beating dynamics which propel the fish. Zebrafish also have pectoral fins, a dorsal fin and an anal fin, and both fins and the body appearance are relevant for signalling and displays [47]. Finally, zebrafish are known to bite [47], but to detect such a behavior would require significantly higher imaging resolution.

While we have focused on pairwise contests between adult male zebrafish, our imaging apparatus and analysis techniques can be readily used to study other social behaviors in zebrafish, or in other swimming animals. One example is aggression in female-female interactions, which are understudied in zebrafish [48, 49]. Another example is courtship, where quantitatively little is known about the behavioral processes that lead to reproductive success [50, 51], though see [52]. Finally, many other fish species engage in pairwise contests, courtship, and other social behaviors. Extending our approach to these systems could address important questions of universality. High-resolution tracking and quantitative analysis of social behavior offers both tremendous potential but also challenges. Single-animal behavioral studies generally rely on egocentric representations of the individual’s pose [11, 12]. However, when studying systems composed of multiple individuals we cannot simply use multiple egocentric coordinate systems, as this omits the relevant spatial configuration of the individuals. Simply put, we need to know both shape of each individuals body, and how they are positioned relative to each other in their environment. A related challenge is the choice of coordinate origin. Do we arbitrarily pick an individual on which to center the allocentric coordinates? And if we do, then how do we combine results from different trials? Here, we used the symmetry-breaking of winner and loser, an effect that will not generally exist in other social contexts.

Advances in imaging and machine vision have enabled high-resolution tracking of animal behavior, though there have been substantially fewer efforts in the settings of social environments containing other animals. But social behavior is universally important in the animal kingdom and beyond, and we hope that our work here provides a useful example of a quantitative approach enabled by new tracking and analysis methods. As we increase the sophistication of our experimental systems and analyses, we can even hope to approach questions traditionally associated with our own human experience, but from a novel direction: one rooted in a complete quantitative language that embraces natural behavior, strives to be free of constraining assumptions, and attempts to infer rather than postulate the behavioral strategies employed.

## METHODS

### Data, Code, and SI Movie Availability

The code for this project can be obtained from the following Github repository: https://github.com/liamshock/Dynamics_of_dominance. A link to download the data for this project, and a link to download the supplementary movies, can be found in the Github README file.

### 3D imaging overview

We constructed a 3D imaging apparatus with an observation tank made of transparent acrylic sheets (thickness of 0.8 cm) assembled to a cuboid structure (inner volume of 40 cm × 40 cm × 44 cm). The imaging volume was 40 cm × 40 cm × 37 cm, with fish-facility produced water (temperature ∼ 35 ^*°*^C, pH ∼ 7, conductivity ∼ 300 uS/cm) filled up to a consistent 36.5 *±* 0.5 cm level across all experiments. The water was then cooled to room temperature, ∼ 27 ^*°*^C. We mounted three USB3.0 cameras (Camera model: CM3-U3-13Y3M-CS, Chameleon3 Mono) in an orthogonal configuration on the camera frame. We positioned the two side cameras ∼ 90 cm away from the front surface of each respective side of the tank, and the top camera ∼ 100 cm above the water surface. While only two cameras are needed for 3D reconstruction, the extra camera was important for reducing occlusions, which are ubiquitous in strongly-interacting conditions. The cameras were controlled by custom-built LabVIEW (National Instruments LabVIEW 2017 SP1 (64-bit)) software installed on a computer (Dell Precision Tower 5810). We streamed images directly into a 4TB Solid State Drive. Each camera could record at up to 150 frames per second (fps) in 8-bit grayscale fullframe (1024 × 1280 pixels). For the dominance dynamics explored here, all experiments were performed at 100 fps for 90 minutes or longer, allowing us to capture a wide range of behaviors, from fast tail beating to the slower dynamics of the dominance decision.

### Imaging illumination

For each camera view, as shown in SI Fig. S1, the observation tank was illuminated with an LED square panel light for backlight illumination, giving a clear visual contrast between the background and the foreground of the zebrafish in the tank. The illumination from respective side panels for each view allowed detection of surface features of the zebrafish, which we used for body-point tracking. A flicker-free LED driver ensured no systematic intensity fluctuation across frames. We placed a white matte-type diffuser sheet of 39.5 cm × 39.5 cm × 0.1 cm on the bottom of the tank to enhance uniformity in lighting and to reduce the reflections on the bottom. We also attached the same diffuser sheet on the two outer sides of the tank in front of the light panels for uniformity in lighting from the sides. With the water in the tank, the light intensity on the exterior sides of the observation tank that were furthest from each respective panel light (with the other two light panels off) was measured an average of 3600 lux, an intensity level that did not noticeably perturb zebrafish behavior due to the lighting condition. As any visual changes of the outside environment can influence the zebrafish behavior, we attached white cardboard to the camera frame (not shown in SI Fig. S1).

### Interior cage

Our apparatus includes a custom-built interior cage, SI Fig. S1(B), placed in the middle of the water-filled tank. The interior cage resolve two important concerns: reflection artifacts that arise when the fish are close to the borders of the tank (the air/water interface and the water/acrylic/air interfaces), and unwanted behavior as zebrafish often interact with their mirror images. The interior cage has two main components: the supporter acrylic frame and the Fluorinated Ethylene Propylene film (also known as the FEP film). The latter is optically transparent in water with a refractive index (*n* ∼ 1.34) very close to water (*n* ∼ 1.33) and high transparency (*>* 90%), both of which reduce reflection artifacts and produce high-quality images. Long-term use of the interior cage is possible due to its strong durability and high chemical resistance to saline water. The design of the acrylic frame provided strong support for the cubic structure of the FEP film.

### Calibration

We quantify fish posture through the automatic identification of multiple bodypoints. To compute the 3D lab coordinates of a fish bodypoint (which must be detected in two or three camera views) we calibrated the imaging volume using a custom-made frame composed of physical beads. We used the centroid positions of 42 beads (∼ 2 mm diameter) as control points. The beads were non-occluding and well-distributed in the image space of all views. The unique positioning of the beads allowed sampling of the whole imaging volume of the tank through 4 rotated configurations, effectively sampling the volume with 168 well-distributed beads control points in the lab space (see SI Fig. S2). Using the image coordinates and measured lab coordinates of the calibration beads, we regressed 5 functions. The function *f*_1_ : (XZ × XY × YZ) → XYZ maps a triplet of image coordinates (one from each view) to a 3D lab coordinate while *f*_2_ : XYZ → (XZ × XY × YZ) performs the opposite. The other three functions each map a pair of image coordinates taken from 2 cameras to an image coordinate in the third, *f*_3_ : (XZ × XY) → YZ with *f*_4_ and *f*_5_ defined similarly by cyclic permutation. To model the non-linearities of the image distortions and Snell’s law, we performed the regression with a polynomial basis expansion. *f*_3,4,5_ were obtained from ridge regression (*α* = 0.01) and a 3rd order polynomial basis expansion of the input image coordinates. *f*_1_ was obtained from ridge regression (*α* = 0.1) and a 2nd order polynomial basis expansion of the input image coordinates. The alpha parameter and polynomial order were determined using a cross-validated grid-search over alpha parameter values and polynomial orders using 3-fold cross validation.

### Animal husbandry protocol

We used 12 adult male zebrafish (*Danio rerio*) of 14-month-old heterogeneous Okinawa wild-type strain (*oki*). We grouped this population into six pairs based on similarities of body size (i.e., in standard length from the tip of the snout to the base of the caudal fin), genetic background and age (i.e., siblings or close relatives bred on the same day). Each pair was housed in a small 1.8 L housing tank with a clear PVC divider, a housing strategy that is typically employed in preparation for zebrafish breeding at the OIST zebrafish facility. The PVC divider ensured that each pair was physically, but neither visually nor chemically, isolated from each other at all times prior to the pair experiment. All zebrafish were maintained according to the standard practices of the zebrafish facility [53].

### Behavior interaction protocol

We began each behavior experiment by using a sterilized net to carefully transfer each fish from their housing tank to the interior cage in the observation tank. With the fish pair now in the observation tank, we enclosed the interior cage and stabilized the cage in the center of the tank. To maintain organism identity, we returned each fish to the same housing tank at the end of the experiment. We started recording just before the fish were placed in the observation tank and we discarded early images, which were distorted due to water ripples. We performed repeated experiments where we matched the same individuals with a resting period of 9 days or longer between contests over six trials. All experiments were performed at 14:00 or later, approximately an hour after which the zebrafish were no longer fed on the same day. Prior to each experiment, we replaced the housing tank water with the circulating water to avoid introducing any particulates of fish skin accumulated through weeks of housing. This step is beneficial not only for image quality (i.e., due to particulates seen in images), but also because fallen fish skins are known to release ‘alarm substance’ that may potentially influence the behavior of the pair at the time of contest [54–56].

### Ethics statement

All experiments were performed in accordance with the Animal Care and Use Program of Okinawa Institute of Science and Technology (OIST) Graduate School, which is based on the Guide for the Care and Use of Laboratory Animals by the National Research Council of the National Academies and has been accredited by the Association for Assessment and Accredition of Laboratory Animal Care (AAALAC International). The protocol was approved by the OIST Institutional Animal Care and Use Committee.

### Dataset

We produced a novel dataset consisting of 22 imaging experiments displaying vigorous interaction activity among six pairs of adult male zebrafish; each recorded in three views at 100 fps for a duration of 90 minutes or longer (up to ∼ 5 hours) in full-frame of 1280 × 1024 pixels, 8-bit grayscale. For each experiment, we saved a set of three uncompressed data in binary RAW format for all views. We transferred the dataset to the OIST computational cluster for a high-quality image data compression using FFmpeg with a constant rate factor of 22 and a “very slow” preset encoding and for storing. We first split the dataset into multiple 1-minute segments, which we then compressed all the segments in parallel, leveraging the computational power of the cluster for processing speed. The output of the individual segments is a set of MP4 files, and we removed the first several minutes of recording to exclude the interior cage adjustment scene from the analysis. Overall, the dataset encompasses ∼ 16 million frames, each frame with 3 views.

### Choice of bodypoints

We track 3 bodypoints on each animal (see Fig. 1). We note that we are measuring 3D bodypoint positions, meaning each bodypoint is a 3D location on the fish body, as opposed to a 2D surface feature. Our first bodypoint, hereafter called the head point, can be thought of as the mouth of the animal (although the mouth itself is not always visible in the images). The second bodypoint, hereafter called the pectoral point, is defined via the pectoral fins. The two pectoral fins are symmetrical on either side of the body of the animal. The pectoral point is defined as the center of mass of the two points which act as the base of the two pectoral fins. This means that the pectoral point is actually placed at an internal point of the fish, with the base of the pectoral fins acting as landmark surface features. From the topdown view of a fish, the pectoral point is seen as the point along the backbone between the bases of the two pectoral fins. The third bodypoint, hereafter called the tail point, is defined as the point on the caudal fin of the fish where the main rigid part of the fish ends, before the flexible end of the caudal fin begins. The tail point is again at a 3D location within the body of the fish. This set of bodypoints provides important behavioral information, including (i) an approximation of the center of mass of the animal (the pectoral point), (ii) an approximation of instantaneous heading of the animal (the unit vector from the pectoral point to head point), and (iii) a measure of the tail beating of the animal (the pectoral point to tail point angle). Hereafter we call this (head, pec & tail) collection of bodypoints, the skeleton.

### Annotation of SLEAP networks training data via 3D GUI

When generating the training data for the 2D SLEAP networks, it was important to enforce 3D consistency in the labelled bodypoint positions. For example, when labelling the image coordinates of the head of a fish in the XZ camera view, the associated labelled image coordinates for the head in the XY and YZ camera views should all represent the same point in physical space. This not only helps with labelling occluded bodypoints which are visible in other camera views, but can be utilized downstream when registering detected bodypoints across camera views (see Methods: Cross-camera registration). To enforce 3D consistency when generating the labelled training data, we built a custom GUI in the JupyerLab environment [57]. The GUI allows an annotator to view all three camera views for a timepoint simultaneously. To label a bodypoint position, the annotator first chooses one of the three camera views to lock in place, meaning the bodypoint image coordinates in this camera view will not be adjusted. When the annotator then adjusted the bodypoint image coordinate in one of the other two camera views, the bodypoint image coordinate in the third camera view is automatically calculated and updated using the image coordinate of the locked camera view, the image coordinate on the adjusted camera view, and the appropriate calibration function (see Methods: Camera calibration). This processes enforces 3D consistency in the labelled points through the regressed calibration functions. The annotator can iteratively refine the image coordinates in each camera view by permuting the locked camera view. In practice, annotators generally locked the camera view where the bodypoint was least visible. Ultimately we gathered a dataset of N=963 training examples for each 2D SLEAP network, taken from N=2 different pairs of fish, using N=5 different annotators. To minimize variability in the placement of bodypoint positions, the 5 annotators first agreed on the working definition of the three bodypoints based on the anatomical features of the fish, and then followed some examples together.

### Cross-camera registration

We define cross-camera registration as the process by which we associate detected 2D skeletons across camera views to get 3D skeletons. The process uses the regressed cross-camera registration functions (*f*_3_, *f*_4_, *f*_5_) to check if bodypoints detected in separate camera views represent the same point in 3D space, and if 2D skeletons detected in separate camera views are observations of the same skeleton in 3D space. Given a 2D image coordinate *a* from camera-A and a 2D image coordinate *b* from camera-B, we say we register *a* and *b* if we associate them as observations of the same 3D point in space from different camera views. We first define a cost of registering single bodypoints *a* and *b* across camera views. The key concept we use is that if *a* and *b* represent the same point in 3D space, then the calibration functions should work reliably. We can compute the point *c* in camera-C that is associated to *a* and *b* by inputting *a* and *b* as arguments into the appropriate function *f*_3,4,5_. If *a* are *b* really are observations of the same point in 3D space, then it should be possible to map from any two of *a, b, c* to the third point with low error. Similarly, if *a* are *b* are not observations of the same point in 3D, then *a, b, c* are not associated with each other via the calibration, and there will be high error in mapping from any two of {*a, b, c*} to the third point. After computing *c*, we compute *f* (*b, c*) = *a*∼ and 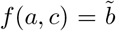 using the appropriate choices of *f* . If *a* are *b* really are observations of the same point in 3D space, ∥*a* − *a*∼∥ and 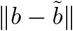 will both be small. Therefore we define the cost of registering *a* and *b* as 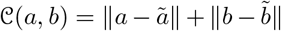. We define the cost of registering two skeletons as the mean of the costs of registering the individual bodypoints. In some cases 2D skeletons may not contain image coordinates for all three bodypoints. We chose to only register skeletons with two or more bodypoints in common. Consider the case where the SLEAP networks have detected two complete 2D skeletons, one for each fish, in each of the three camera views. This example represents the maximum possible available information for the cross-camera registration step. To perform the cross-camera registration, we first measure the skeleton registration costs for all of the 4 permutations possible when matching the skeletons across all three camera views. With the skeletons now assigned across camera views, the final step is to get the 3D bodypoint positions. At this point we could input the image coordinates from all three camera views into the regression function *f*_1_ to get 3D bodypoint positions. But in practice we found that we obtained smoother trajectories by first identifying the pair of cameras with the lowest cost of registering both skeletons, defined as the sum of the skeleton registration costs, and using these 2D image coordinates and the appropriate calibration function *f*_3,4,5_ to derive the 2D image coordinates in the third camera view. Then using the detected bodypoints from the 2 registration cameras and the computed image coordinates from the third camera view, we compute the 3D bodypoint positions using *f*_1_. The motivation for why this provides smoother trajectories is that by following this process we are always inputting image coordinates into *f*_1_ that are most representative of the same point in 3D space as seen by our calibration functions. The cross camera registration process becomes more complicated when we are missing 2D image coordinate data from the SLEAP networks. This can be due to fish-fish occlusions, fish self-occlusions by orientating themselves along the optical axis of the cameras (particularly common in the two side camera views), or occlusions caused by acrylic structure which mounts the interior cage. The process as described in the above paragraph relies on choosing the minimum of a set of possible registration possibilities, and relying on the permutation with the lowest cost being the correct registration. But consider the example of obtaining only a 2D skeleton for fish-1 in camera-A and a 2D skeleton for fish-2 in camera-B. In this case, we have only one possible registration, and it is incorrect. Motivated by such situations, we introduced a threshold on skeleton registration costs. In frames where we have two detected skeletons in at least two camera views, we can readily identify the correct registration without using a threshold, simply by choosing the permutation with the lowest cost. Then by examining the distribution of registration costs estimated over all such frames, we determined a threshold for correct registrations as reg thresh = 10 (see SI Fig. S3). We then use this threshold to accept or reject registrations from frames without two detections in at least two camera views. Another complication is that of missing bodypoints. When computing the cost of registering skeletons across camera views, if even one of the skeletons has a missing bodypoint (no SLEAP information for that point) then the cost of registering that bodypoint is undefined. We made the decision to only register skeletons that had at least two of the three bodypoints detected.

### Post-processing idtracker.ai trajectories

We apply idtracker.ai [32] to the top-down XY camera view, resulting in identity-registered centroid trajectories of both individuals. We observed infrequent discontinuities in the idtracker.ai trajectories. Before using the idtracker.ai data to register identities for the SLEAP data, we post-processed the idtracker.ai data. For all fish from all recordings, we computed the absolute value of the acceleration of the centroid timeseries, and examined the distribution (see SI Fig. S3). We set a threshold value of 3.3, and removed data from the timeseries 6 frames either side of frames which exceeded the acceleration threshold.

### Identity registration

We apply idtracker.ai [32] to the top-down XY camera view, resulting in identity-registered centroid trajectories of both individuals, with gaps when images of the animals overlap. We use these centroid trajectories to assign identity to each 3D skeleton. We identify all frames where, for both fish, we have XY image coordinates of the pectoral point, as well as idtracker.ai XY centroid image coordinates. Using the pectoral point as an approximation to the centroid, we register the idtracker.ai centroids to the XY pectoral point image coordinates using the Hungarian sorting algorithm [33], with the euclidean distance between 2D image coordinates as the cost. For the majority of frames (those where fish images are not overlapping) we use this process to assign global identity to 3D skeletons. In our dataset, the experiment-mean percentage of frames that contained idtracker.ai centroids for both individuals was ∼86%. More then 99% of frames with idtracker.ai centroids for both individuals also contained XY pectoral points for both individuals. The final step of the identity registration process is to match idtracker.ai centroids to SLEAP skeletons in frames where we do not have two idtracker.ai centroids and two SLEAP skeletons. Similarly to the above section on cross-camera registrations, we examine the distribution of idtracker.ai-to-SLEAP registration costs in frames where we can use the Hungarian algorithm, and hence determine a threshold for correct registrations as reg thresh = 12.5 (see SI Fig. S3). We use this threshold to assign identities wherever possible in the frames without two idtracker.ai centroids and two SLEAP skeletons.

### Identity propagation via 3D bodypoints

For frames in which identity cannot be directly assigned as above (resulting from image occlusion), we propagate identity using a frame-to-frame tracking of 3D skeletons. First we identify contiguous temporal segments of the experiments where we could not register identities to 3D skeletons using idtracker.ai results. For each of these segments, we have available 3D skeletons with identity, just before the start of the segment and just after the end of the segment. We attempt to propagate identity through these segments using a frame-to-frame tracking of 3D skeletons. Given two 3D skeletons in frame *t* with global identity, and two 3D skeletons in frame *t*+1 without global identity, we propagate identity from frame *t* to frame *t* + 1 by registering 3D skeletons in frame *t* to frame *t* + 1 using the Hungarian sorting algorithm, with the sum of the euclidean distances between bodypoints as the cost function. Since we know the positions of the 3D skeletons with identity just after the end of the segment (from idtracker.ai), we can determine if the attempted tracking of the segment was successful, i.e., did we end up with the two individuals in the correct locations, or did we swap identities during the tracking of the segment and end up with the two individuals in the wrong locations. Segments which were successfully tracked (correct locations for individuals after the segment) were included in the dataset. Segments which were unsuccessfully tracked were removed from the dataset to avoid, as much as possible, the final data containing 3D skeletons with incorrect identities. See Table S1 for statistics on the quality of the tracking.

### Trajectory post-processing

Occasionally we will produce a 3D skeleton which is unnaturally large. This can be caused by amplification of network noise through the calibration functions, or mis-registrations across camera views (which can be the result of errors in the 2D skeleton output of SLEAP, such as for example both contestants being assigned the same head position). Across the recording, we measure the mean distance between the head and pectoral points, and the pectoral and tail points. We remove from the dataset any 3D skeletons where the head-pec distance is greater than 1.2cm or the pec-tail distance is greater than 2.5cm (see SI Fig. S3). Finally we interpolate and filter the 3D bodypoint trajectories, working individually across each spatial dimension. We fill-in gaps to a maximum size of 5 frames using a first order polynomial function. We then filter the trajectories using a Savitzky–Golay filter with 2nd order polynomial over a 9 frame window.

### Identification of fight bouts

We identify fight bouts by combining intuition developed from ethological measures [28], with a new, quantitative behavioral clustering, Fig. 3. We cluster time-windowed probability distributions of our configuration variables. When identifying fight bouts we want our method to be agnostic to which fish ultimately wins the contest. We do not want the labelling of fish-1 and fish-2 to affect the outcome of the clustering process. To this end, we do not work directly with the state variables (*θ*_1_, *θ*_2_). We use the symmetric function *g*(*θ*_1_, *θ*_2_) = *g*(*θ*_2_, *θ*_1_) = (*θ*_1_ + *θ*_2_, |*θ*_1_ − *θ*_2_|), and we cluster time-windowed probability distributions of (*D*_*P P*_, *θ*_1_ + *θ*_2_, |*θ*_1_ − *θ*_2_|). To discretize the (*D*_*P P*_, *θ*_1_ + *θ*_2_, |*θ*_1_ − *θ*_2_|) variables, we used *N* = 20 partitions for each variable, for a total of 20^3^ = 8000 microstates. We set the lower limit for the *D*_*P P*_ discretization as 0, and the upper limit as 20. We set the limits for the (*θ*_1_ + *θ*_2_) discretization as −*π* and *π* respectively, and we set the limits for the |*θ*_1_ − *θ*_2_| discretization as 0 and *π* respectively, so the upper and lower limits for the discretization of the variables span the domains of the variables. The time windows are 1min in duration, where the duration is chosen to be an order of magnitude larger than the correlation time of the input variables. We compute the (*D*_*P P*_, *θ*_1_ + *θ*_2_, |*θ*_1_ − *θ*_2_|) probability distribution in each window. We define the similarity between each pair of sampled distributions using the Jensen-Shannon distance

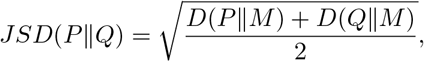

where *P* and *Q* are probability distributions, *M* is the point-wise mean of *P* and *Q*, and *D* is the KullbackLeibler divergence. In the first pass at fight bout identification (before we refine the boundaries), we use 1min time windows with 30sec overlap, using data from all 22 experiments, for a total of 5308 probability vectors. We then use hierarchical clustering (python scipy.clustering package [58]) to cluster the probability distributions, where the dendrogram reveals a clear branch associated with fight bouts (see SI Fig. S7). We then merge fight bouts which are separated by less than 2.5min, and remove fight bouts smaller than 2.5min. We then refine the boundaries of the identified fight bouts (second pass). In the first pass the overlap of adjacent windows is 30sec. For the second pass, we create a set of windows before and after the start and end of the boundaries from the first pass, with 58sec overlap, with sufficiently many time windows to extend 30sec before and after the first pass fight boundaries. We then label these new time windows as fight windows or non fight windows by re-clustering them. In each new time window, we compute the associated probability distribution, find the nearest neighbor in the originally clustered set of probability distributions, and assign the new time window the label of its nearest neighbor. We set the refined start time of each fight bout as the center of the first time window in the refining set of windows for the bout which was assigned the label of fight. We set the refined stop time of each fight bout as the center of the last time window in the refining set of windows for the bout which was assigned the label of fight.

### Winner/loser assignment

To determine the winner/loser assignment in each imaging experiment we consider the epoch stretching from the end of the last fight bout to the end of the imaging experiment. In this epoch we estimate the distribution of *θ*_1_ and *θ*_2_. For the example shown in Fig. 3 there is a clear asymmetry: one animal is pointing much more frequently towards the other. We identify this animal as the winning fish. To generalize from this example, across all experiments, we compute the summed probability around the *θ* = 0 peak, 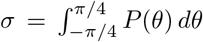, and we find a bimodal distribution, SI Fig. S9. We assign winner/loser labels based on whether *σ* falls in one peak or the other.

### Transition dynamics in the joint-animal “state space” spanned by (*D*_*PP*_, *θ*_*W*_, *θ*_*L*_)

We characterize two-fish dynamics through transitions between microstates of the relative distance and orientation angles. To discretize the configuration variables, we used *N* = 20 partitions for each variable, for a total of 20^3^ = 8000 microstates. The upper and lower limits for the *θ* discretization were set as − *π* and *π* respectively to span the entire domain of the *θ* variables. The domain of *D*_*P P*_ ranges from 0cm to ∼ 52cm (the distance between opposite corners of the interior cage). We set the lower limit for the *D*_*P P*_ discretization as 0cm, and the upper limit as 20cm. This upper limit is set large enough to avoid artificially truncating fight maneuvers which occur at close distance. We estimate a transition matrix *T*_*ij*_ by computing the transition probability among all microstates within a timescale *τ*, which is a free parameter. We fix *τ* by examining the emergence of macrostates through Infomap clustering [35], as described below.

### Finding two-fish maneuvers by coarse-graining the transition dynamics using Infomap community detection

To study contest dynamics encoded in the transition matrix *T*_*ij*_, we perform community detection using a Python igraph [59] implementation of the Infomap algorithm [35]. We first convert *T*_*ij*_ to a weighted directed graph, removing any null states. We then execute the Infomap algorithm, which returns a set of transition matrix network communities, collections of microstates which are clustered together by the dynamics of *T*_*ij*_, and which we denote as maneuvers. We set trials=500, which sets the number random walker placements. All other parameters of this implementation match the default input parameters of the original multi-level partitioning Infomap algorithm [35].

### Determining the transition time *τ*

To identify an appropriate choice of *τ* we examine the structure of the Infomap clustering of *T*_*ij*_ as a function of *τ*, SI Fig. S10 (A), focusing on the volume fraction of the largest cluster. As we approach *τ* = 12 frames, we find a sudden large jump, which occurs from merging all of the close distance clusters. We choose *τ* = 10 frames to simplify the number of emergent communities, while simultaneously avoiding this sudden loss of resolution.

### Maneuver descriptions

Here we provide a description for each of the *N* = 10 fight maneuvers motivating the descriptive label we attached to each. **Attack** We ascribe the label of “attack” to one set of two clusters (see Fig. 5 & SI Fig. S11, as consistent with previous descriptions [28]. In these states, the attacker is positioned behind the attackee, with the attacker pointing towards the attackee and the attackee pointing away from the attacker, at close distance. Regardless of whether the attacker makes physical contact during these cluster dynamics, the attack is still in an advantageous position relative to the attackee. **Approach** We ascribe the label of “approach” to one set of two clusters (see SI Fig. S11). The relative orientations of the “approach” maneuvers are similar to that of “attack” maneuvers, but “approach” maneuvers occur at further distance than “attacks”. Activations of “approach” maneuvers are consistent with previous descriptions of chasing [28], and are often followed by an “attack” as the distance between the animals reduces. **Close circling** In the cluster we label as “close circling”, the two animals are in an anti-parallel configuration at close distance (see SI Fig. S11). The pair tend to spin around each other during “close circling” (large 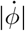). Spinning in an antiparallel alignment at close distance motivated the label of “circling”. We remark that the animals do not have to make a full rotation, or multiple rotations, during a “circling” maneuver activation. **Display** In the cluster we label as “display”, the two animals are in a parallel configuration at close distance (see SI Fig. S11). Close distance parallel alignments are consistent with previous descriptions of lateral displays [47]. **Direct Towards** In the cluster we label as “direct towards”, both animals are pointing directly towards the other (see SI Fig. S9). The *D*_*P P*_ range spanned by this maneuvers runs from 1cm to 9cm. Activations of this maneuver facilitate the pair closing the distance between them. However, both fish do not have to be actively swimming towards each other during this maneuver. One fish can be static in the tank pointing towards the opponent, while the other fish swims directly toward the first. **Direct Away** In the cluster we label as “direct away”, both animals are pointing directly away each other (see SI Fig. S11). The *D*_*P P*_ range spanned by this maneuvers runs from 3cm to 11cm. Activations of this maneuver facilitate the pair moving directly away from each other. **Middle Circling** The cluster we label as “middle circling” has similar structure in the relative orientations as “circling”, but occurs at further distance (see SI Fig. S11). It also has a similarly large 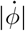 indicating spinning of the system. We use the label “middle circling” as a shortened version of “middle-distance circling”. **Middle Parallel** The cluster we label as “middle parallel” has the pair in a parallel configuration at a *D*_*P P*_ range of 6cm to 12cm. In this case we do not use a term such as “middle-distance display”, as previous descriptions of [47] tend to have the contestants at close to each other. We remark here that this cluster was the last cluster to be included in the set of *N* = 16 clusters which span fight bouts, and has a very low probability of occurring during fight bouts.

## ACKNOWLEDGEMENTS

We thank Keita Kamino for comments on the manuscript. This work was supported by grant RGP0055 from the Human Frontiers Science Program, and by funds from OIST Graduate University (TI, IM, GJS) and Vrije Universiteit Amsterdam (GJS). We thank all members of the Developmental Neurobiology Unit at OIST for general zebrafish maintenance support, and especially Yuko Nishiwaki and Luis Carretero for insightful discussions and for providing the zebrafish. We also thank Daniel Lee for early prototype of the imaging system. We are grateful for the help and support provided by the Engineering and Scientific Computing & Data Analysis sections of Research Support Division at OIST.

## SUPPLEMENTARY MATERIAL

**FIG. S1.**
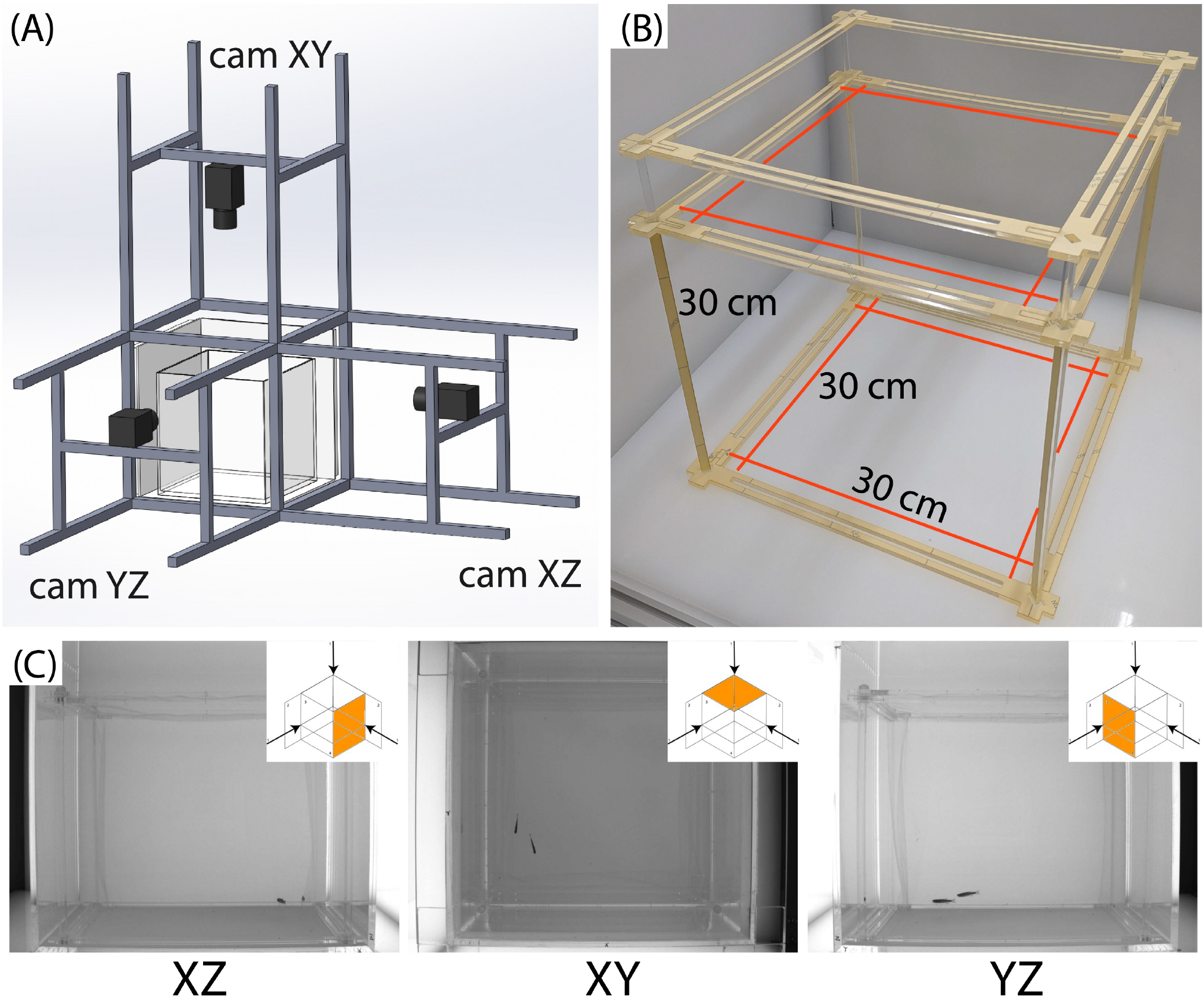
Experimental apparatus: (A) Schematic of the imaging apparatus. Three USB3.0 cameras (Camera model: CM3-U3-13Y3M-CS, Chameleon3 Mono) are mounted in a 3-orthogonal configuration on the aluminum camera frame, with the observation tank (inner dimensions of 40 cm × 40 cm × 44 cm, acrylic thickness of 0.8 cm) in the center of the frame. Two cameras are mounted horizontally ∼ 90 cm away from the respective tank front for the two orthogonal side view imaging, and the other camera is mounted vertically ∼ 100 cm away from the water surface to image the top view of the tank. To minimize any external visual influences on the zebrafish in the tank, white cardboards are attached to the aluminum camera frame to enclose the whole imaging system (not shown). (B) The interior cage acrylic frame (including the top cover frame) is shown before attaching the FEP film and removing the yellow papers on the acylic material. The red lines are drawn to show the placement of the fishing lines for structuring the FEP film into a cube of dimensions 30 cm × 30 cm × 30 cm. (C) Images from the three views (from left to right): XZ (side view), XY (top view), and YZ (side view). These X, Y, and/or Z are marked at a corner of the tank for each view to avoid any confusion, especially between the two side views.

**FIG. S2.**
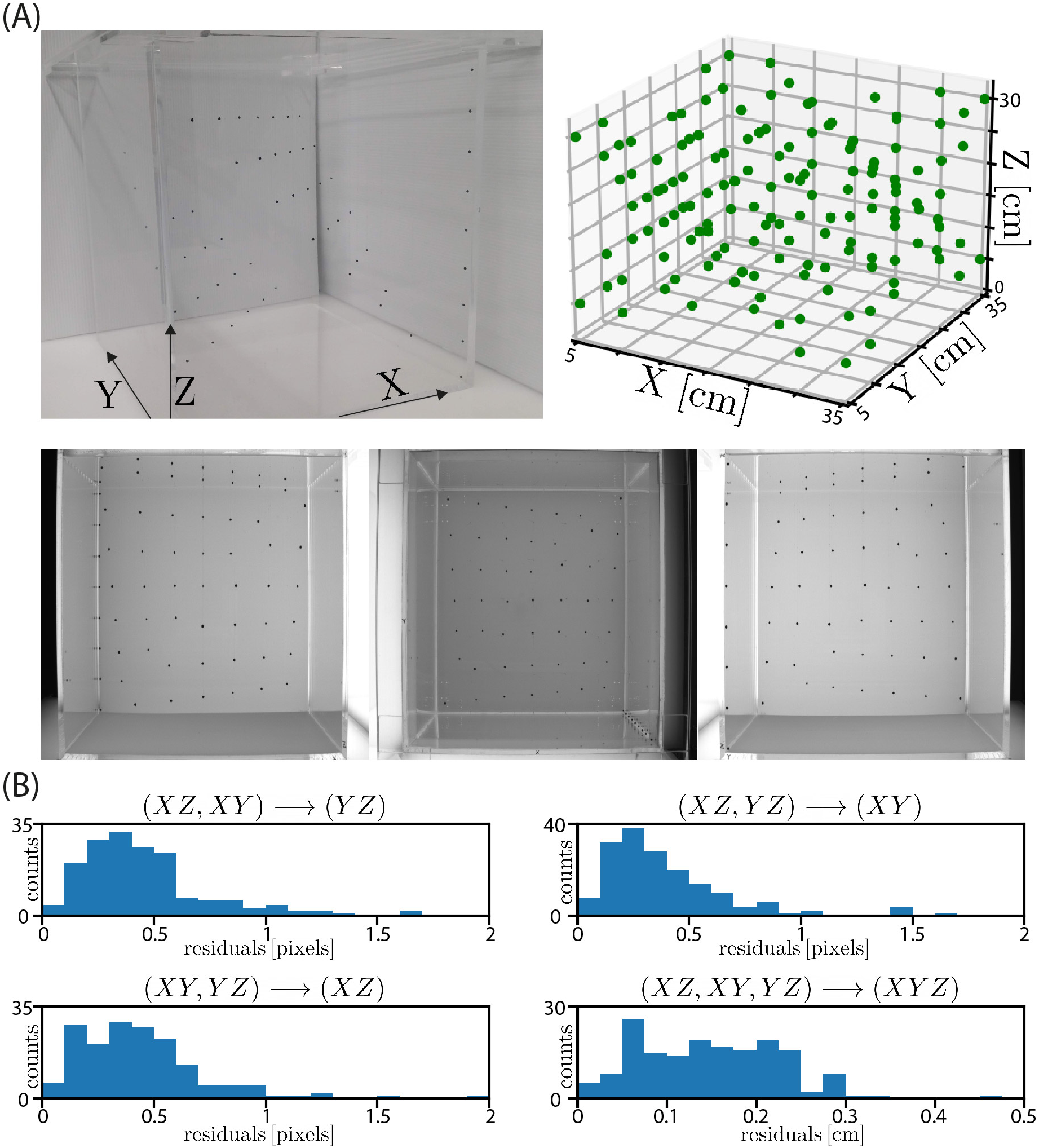
Calibration: To make correspondences between the lab space and the image space of each of the three views, a bead calibration structure was designed to model the observation tank volume. The structure was imaged at 4 rotated configurations to sample the whole volume of the observation tank. The calibration structure is designed such that beads do not occlude each other when viewed through the three cameras in any of the four rotations. (A, top-left) A photograph of the bead calibration structure. The diameter of each bead was approximately ∼ 2 mm. (A, top-right) Schematic of the collection of 168 calibration beads gathered from all four rotations of the calibration structure. To build the calibration models we ultimately only use beads positioned within the volume of the interior cage (SI Fig. S1). The top 7 beads (and their reflected beads directly above) in the XZ and YZ images and 9 edge beads were not used. Effectively, we sampled the whole volume with 168 beads through 4 rotations. (A, bottom) Images from the three views in the presence of the calibration structure immersed into the observation tank in one of the 4 rotations. (B) We regressed four calibration models. Three models take image coordinates from a pair of camera views and return the associated image coordinates in the third camera view. The last model takes an image coordinate from all three camera views and returns the associated 3D position in the tank. We show here the residuals of the calibration models. The residuals of the models mapping two camera views to the third camera view were generally less than 1 pixel. The residuals of the 3D model were generally less than 2mm. Building the calibration models using a polynomial basis expansion of the input coordinates resulted in a smooth mapping of the 3D volume.

**FIG. S3.**
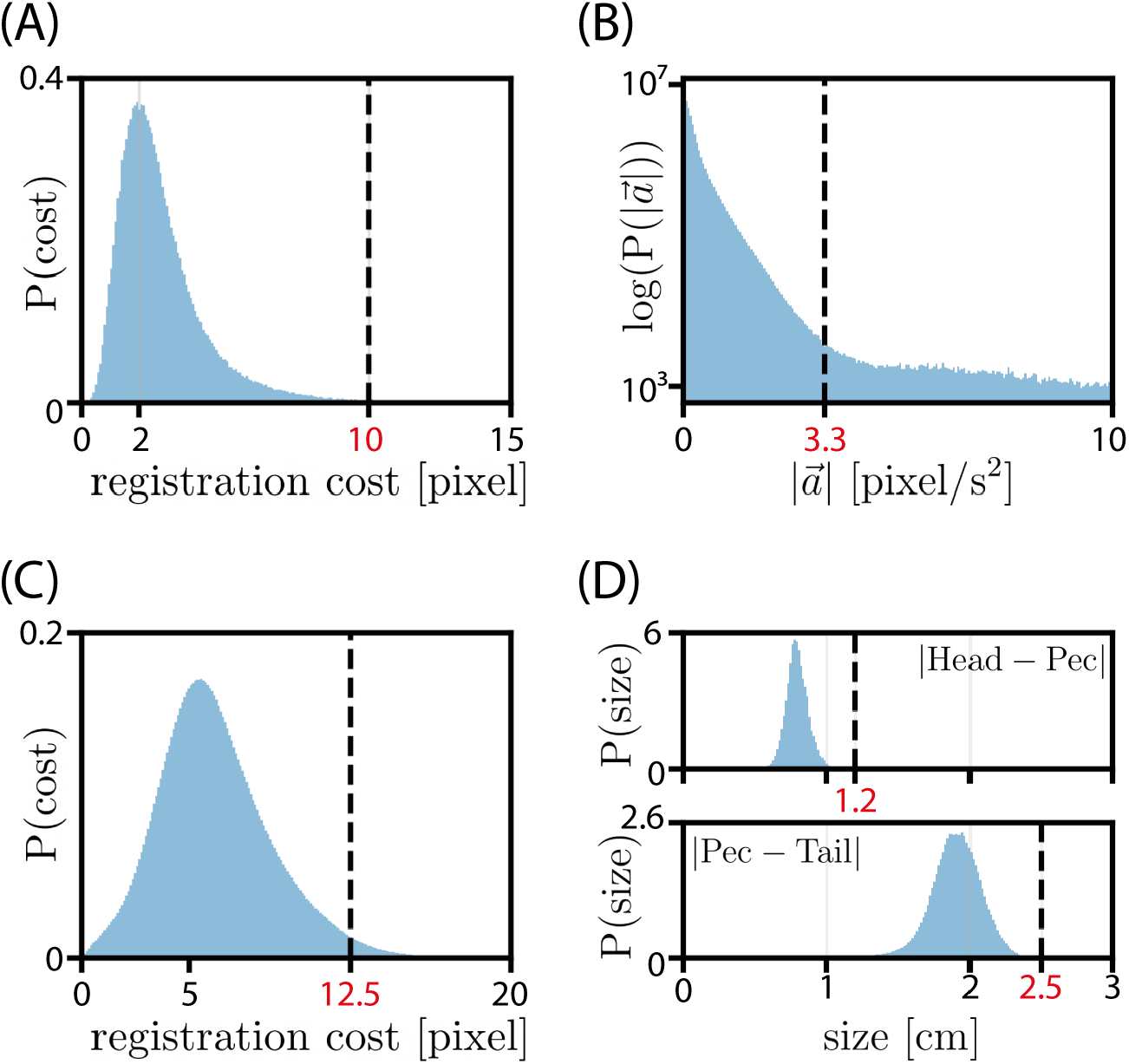
Threshold acceptance values for tracking. (A) We show the distribution of cross-camera registration costs for frames where we know the registrations are correct (frames with two 2D skeletons detected in at least two camera views, see Methods). We set the acceptance threshold at *registration thresh* = 10. (B) We show the distribution of the absolute value of the acceleration of the idtracker.ai centroids. We set the *idtracker acceleration thresh* = 3.3. (C) We show the distribution of costs for assigning idtracker.ai centroids to SLEAP pec-points. We set the *idtracker to sleap thresh* = 12.5. (D-top) We show the distribution of head-pec distances for individuals during all recordings and set a maximum size threshold of *head pec thresh* = 1.2 cm. (D-bottom) We plot the distribution of pec-tail distances for individuals during all recordings. We set a maximum size threshold of *pec tail thresh* = 2.5 cm.

**TABLE I.** Tracking accuracy summary statistics. In this table we present data illustrating the quality of the tracking of the experiments before post-processing of trajectories. In the first column we list the names of the experiments, which contain the experiment start-time in year-month-day hour-min-sec format. In the second column we list the pair index of the contestants. In the third column we list the number of frames in the recordings of the experiments. In the forth column we list the identity accuracy estimate from idtracker.ai. In the fifth column we list the percentage of frames in the recording where we have no available 3D bodypoint information. In the sixth column we list the mean size in frames of temporally contiguous regions of the recordings where we have no available 3D bodypoint information. In the seventh column we list the 99%-quantile, in frames, of temporally contiguous regions of the recordings where we have no available 3D bodypoint information, showing almost all missing data regions are shorter than 0.5 seconds. In summary, we have occasional small gaps in the trajectory data on the order of a few frames which are interpolated over during post-processing, and we have some larger infrequent gaps in the trajectory data which we do not interpolate over and instead leave as missing data. Of the 22 experiments recorded and tracked, 18 were used for subsequent analysis. Two experiments were removed from subsequent analysis because we could not reliably determine the winner and loser. Another two experiments were removed because the contestants in these trials showed a strong dominance relationship from the beginning of the recording, leaving us unable to probe the emergence of the dominance relationship.

| Experiment name | Pair | # frames | idTracker acc [%] | NaN frames [%] | mean NaN-gap [frames] | 99% quantile NaN-gap [frames] |
| --- | --- | --- | --- | --- | --- | --- |
| FishTank20200127_143538 | 2 | 971756 | 0.996 | 5.56 | 3.81 | 45 |
| FishTank20200129_140656 | 3 | 769848 | 0.994 | 6.15 | 3.80 | 40 |
| FishTank20200130_153857 | 4 | 501943 | 0.998 | 3.06 | 2.92 | 30 |
| FishTank20200130_181614 | 5 | 517000 | 0.998 | 2.62 | 2.71 | 25 |
| FishTank20200207_161445<sup>†</sup> | 6 | 463997 | 0.999 | 6.56 | 3.48 | 35 |
| FishTank20200213_154940 | 2 | 730119 | 0.997 | 4.72 | 3.55 | 41 |
| FishTank20200214_153519 | 4 | 495610 | 0.996 | 1.11 | 2.41 | 16 |
| FishTank20200217_160052 | 5 | 601226 | 0.999 | 1.89 | 2.70 | 26 |
| FishTank20200218_153008 | 3 | 537880 | 0.999 | 4.05 | 2.98 | 33 |
| FishTank20200316_163320<sup>†</sup> | 6 | 1032353 | 0.999 | 1.71 | 2.53 | 19 |
| FishTank20200327_154737 | 4 | 561010 | 0.995 | 1.21 | 2.77 | 28 |
| FishTank20200330_161100 | 5 | 556428 | 0.999 | 2.45 | 3.31 | 37 |
| FishTank20200331_162136 | 3 | 746434 | 0.998 | 4.46 | 3.19 | 34 |
| FishTank20200520_152810 | 2 | 1248577 | 0.990 | 2.34 | 2.66 | 22 |
| FishTank20200521_154541 | 4 | 596607 | 0.993 | 1.35 | 3.27 | 36 |
| FishTank20200525_161602 | 3 | 588962 | 0.999 | 6.77 | 3.64 | 38 |
| FishTank20200526_160100<sup>‡</sup> | 1 | 689100 | 0.998 | 5.77 | 3.81 | 44 |
| FishTank20200527_152401 | 6 | 497635 | 0.999 | 1.64 | 2.45 | 18 |
| FishTank20200824_151740 | 2 | 1695643 | 0.990 | 3.19 | 2.97 | 31 |
| FishTank20200828_155504 | 5 | 798382 | 0.998 | 1.75 | 2.69 | 21 |
| FishTank20200902_160124 | 3 | 717814 | 0.995 | 4.47 | 3.42 | 41 |
| FishTank20200903_160946<sup>‡</sup> | 1 | 726000 | 0.997 | 7.27 | 3.16 | 35 |

<sup>†</sup> These experiments did not have a clear winner and loser.
<sup>‡</sup> These experiments displayed dynamics indicative of a dominance relationship from the start of the recording.

**FIG. S4.**
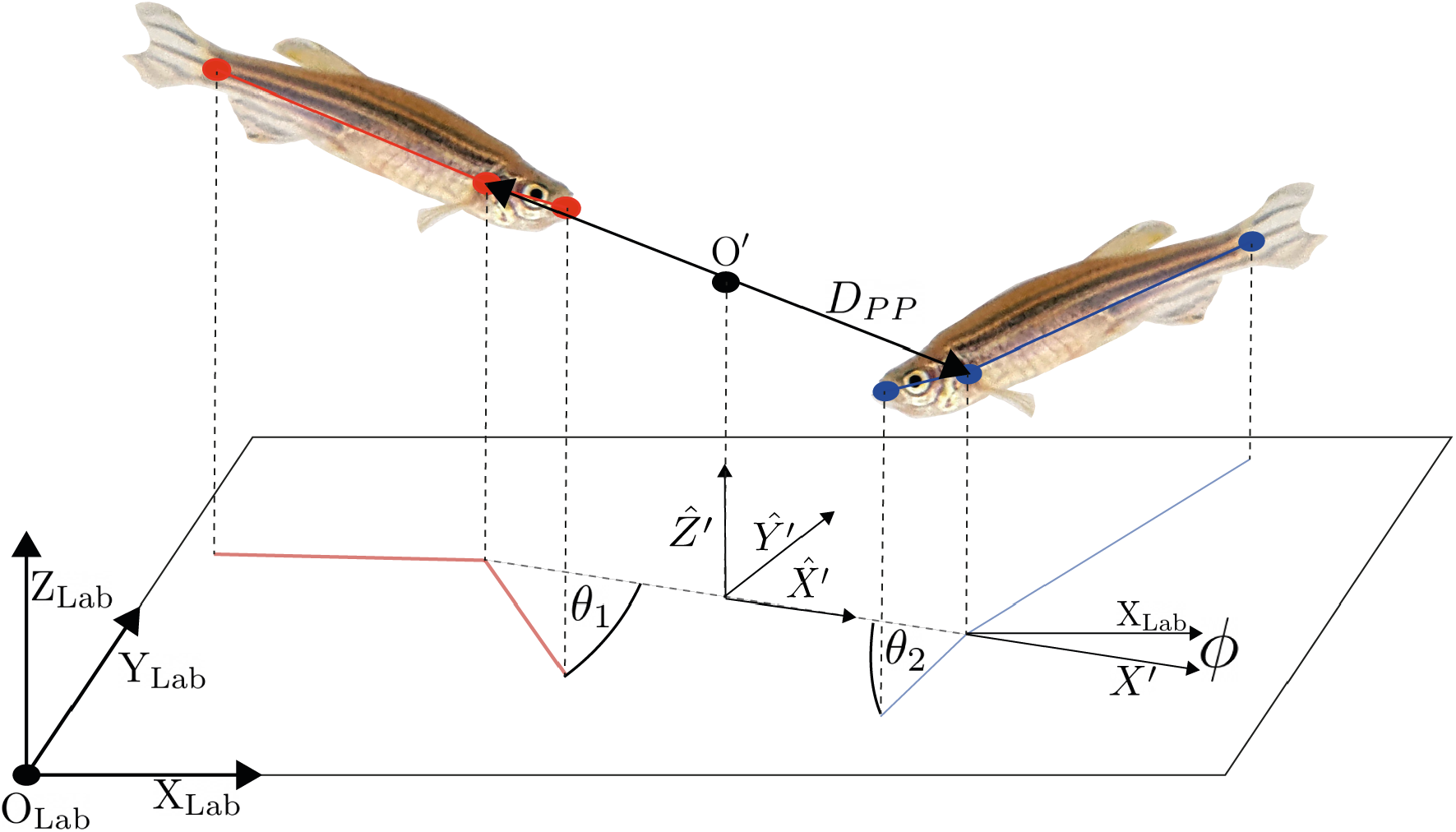
Schematic of the co-rotating coordinate system. The output of the tracking pipeline is the 3D positions of the 3 bodypoints of both fish in laboratory coordinates. Motivated by our desire for a translationally invariant and XY rotationally invariant representation of the fish system, we performed a coordinate transformation to a co-rotating frame. Formally, for each timepoint of the recordings, we moved from writing the 3D bodypoints in the lab coordinate system with basis vectors {*X*_*Lab*_, *Y*_*Lab*_, *Z*_*Lab*_} and origin *O*_*Lab*_, to writing the 3D bodypoints in the co-rotating coordinate system with basis vectors {*X*^*′*^, *Y* ^*′*^, *Z*} and origin *O*^*′*^. Defining the new origin *O*^*′*^ introduces translational invariance. We defined *O*^*′*^ as the center of mass of the “Pec” points of the pair. Gravity acting in *Z*_*Lab*_ breaks the 3D symmetry in the tank, and hence we chose to set *Z*_*Lab*_ = *Z*^*′*^. To introduce XY rotational symmetry, we define *X*^*′*^ as the *X*_*Lab*_ and *Y*_*Lab*_ components of the vector running from the “Pec” point of the winner to the “Pec” point of the loser. We set *Y* ^*′*^ to complete the right-handed basis set {*X*^*′*^, *Y* ^*′*^, *Z*^*′*^}. Setting the new origin *O*^*′*^ removes 3 degrees of freedom from the original 18D Lab coordinate data, and setting *X*^*′*^ removes one further degree of freedom, providing the 14D data on which we perform PCA in Fig. 2(A and B). For clarity, to directly account for the dimensionality of the 14D system, both “Tail” points and both “Head” points have 3 DOF each for a total of 12 DOF. The two “Pec” points have zero *Y* ^*′*^ components by construction, and they have identical magnitude but sign flipped *X*^*′*^ and *Z*^*′*^ components, for 2 DOF of freedom. To define the XY rotational speed of the system 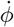, we define *ϕ* as the angle between *X*_*Lab*_ and *X*^*′*^ and take the time derivative.

**FIG. S5.**
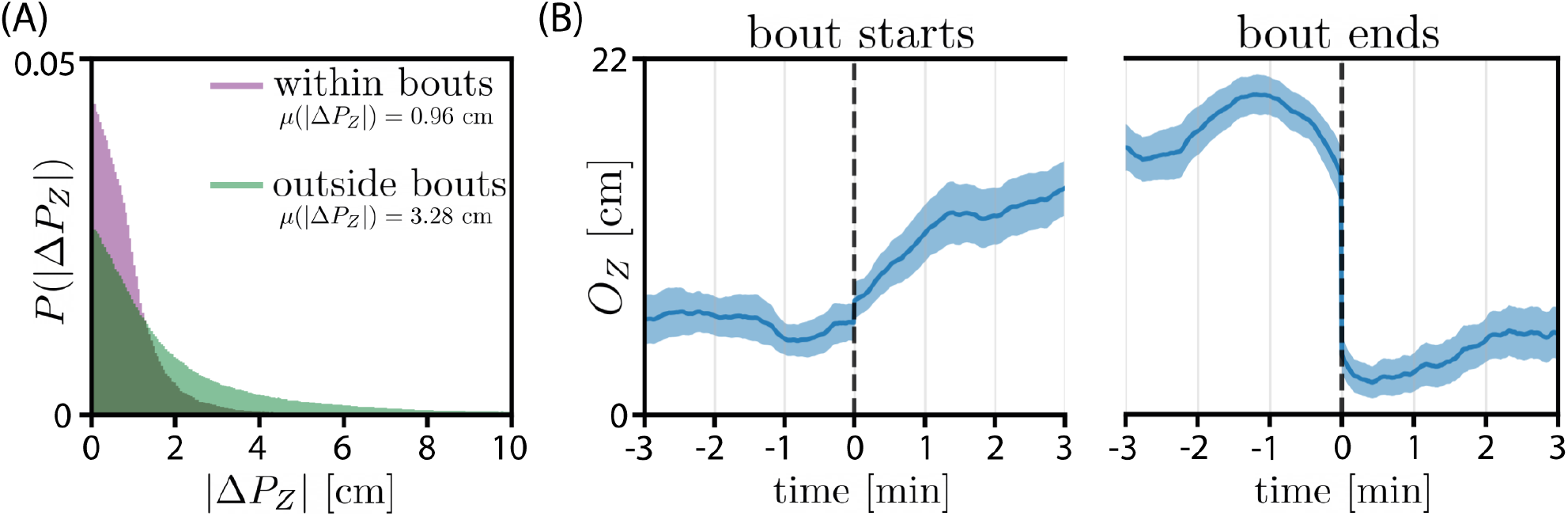
During fight bouts fish are generally close in vertical separation *Z*, suggesting that fighting is a planar activity. The plane tends to move upward in Z as fights progress. (A) We show *P* (|Δ*Z*|) = |*P*_*Z*,1_ − *P*_*Z*,2_|, the absolute value of the difference of *Z* coordinates of the “Pec”“points of the pair, during fight bouts and outside of fight bouts. The mean value of |Δ*Z*| during fight bouts is ∼1 body-height. (B) We show *O*_*Z*_, the *Z*-component of the co-rotating frame origin, aligned to the start (B-left) and end (B-right) of fight bouts. We calculate the mean value of *O*_*Z*_ in 60-sec running windows with 59-sec overlap. Plotted is the mean (dark line) and standard deviation (shaded region) from fight bouts longer than 6-minutes. As fight bouts progress the pair ascends in the tank. At the end of fight bouts the pair abruptly descends toward the bottom of the tank. Taken together, (B-left) and (B-right) motivate a description of fighting as occurring in Z level planes which ascend upward over time during contests.

**FIG. S6.**
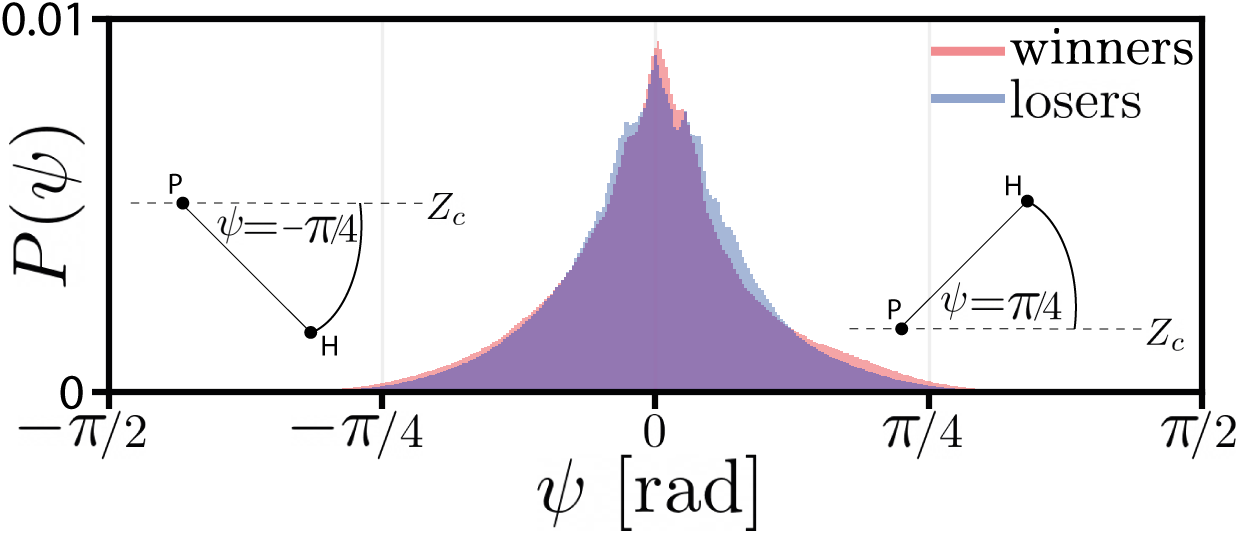
The distribution of the fish pitch angle *ψ* is strongly peaked around *ψ* = 0. We define the pitch angle *ψ* as the angle between the pec-head vector and the Z-level-plane at the pec point. Shown in this panel is the distribution of *ψ* for winners and losers across all experiments. The distribution is peaked around *ψ* = 0, which represents no relative Z displacement between the pec and head points. The pitch angle rarely extends beyond *ψ* = *± π/*4. We use this observation, combined with the small amount of variance explained by the pitch PCA modes (Fig. 2), as motivation for excluding the *ψ* angles from our configuration space.

**FIG. S7.**
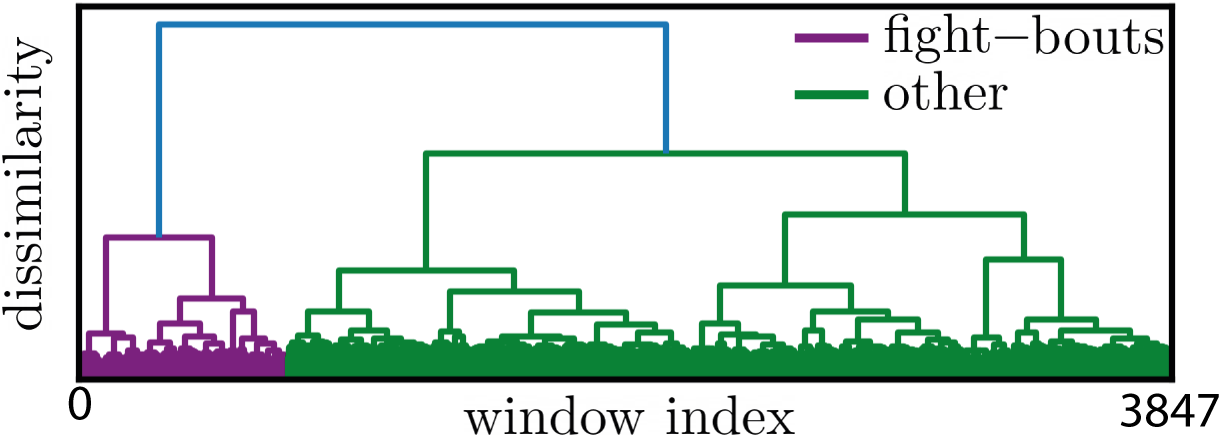
Fight bouts emerge from hierarchical clustering of the 3D distribution of anonymized state variables. Initially we have the state variables (*D*_*PP*_, *θ*_1_, *θ*_2_) which require a choice of which contestant is 1 and which contestant is 2. We apply the symmetric function *f* (*θ*_1_, *θ*_2_) = *f* (*θ*_2_, *θ*_1_) = (*θ*_1_ + *θ*_2_, |*θ*_1_ − *θ*_2_|) = (*θ*_*a*_, *θ*_*b*_) which is invariant under the contestant labelling choice. In 1-min running windows with no overlap, we compute the *P* (*D*_*P P*_, *θ*_*a*_, *θ*_*b*_), for a total of 5308 probability vectors. Using the Jensen-Shannon-distance as a distance metric, we hierarchically cluster the 5308 probability vectors. We show the dendrogram from the hierarchical clustering with a K=2 partitioning. The *D*_*PP*_, *θ*_1_, *θ*_2_ distributions for these clusters can be seen in Fig. 3(B).

**FIG. S8.**
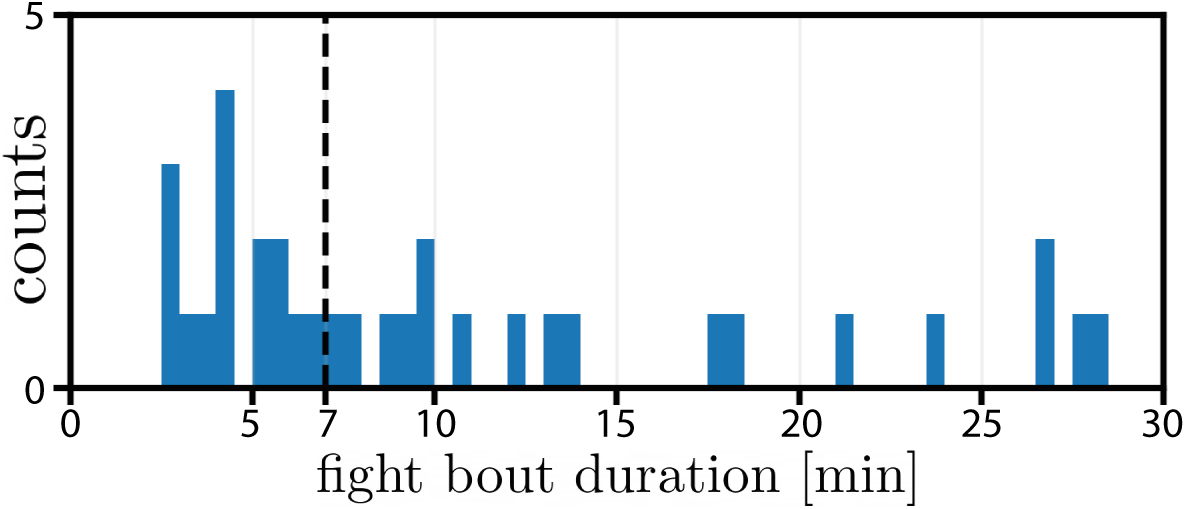
The distribution of fight bout durations. We plot the distribution of fight bout durations computed from fighting epochs identified from our fight bout detector on the N=18 experiment dataset. The vertical black dashed-line at 7 minutes shows the threshold on fight duration used to identify the set of fight bouts used in Fig 6 (B). For the 18 experiments, we identified a total of 33 fight bouts. The mean bout duration was 11 min. 18 fight bouts lasted longer than 7 minutes.

**FIG. S9.**
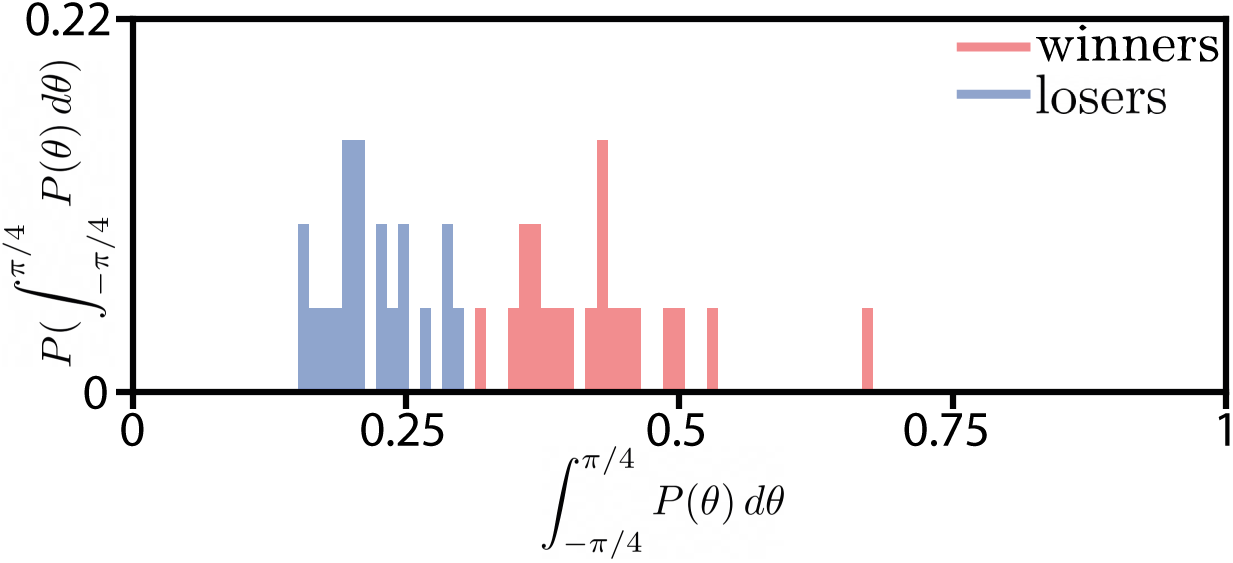
After fight bouts have ended, winner and loser robustly visit different orientation states. To identify the winner/loser we focus on the asymmetry which emerges between contestants after fight bouts have concluded. In Fig. 3(Bbottom) we illustrate the peak centered at *θ* = 0. After the last fight bout in each experiment, this peak is large for the contestant declared the winner and is small for the contestant declared the loser. To quantify the size of this peak, we measure the *θ* probability from − *π/*4 to *π/*4. In each trial 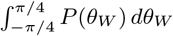 and 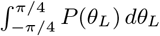 are clearly distinct. To illustrate this point, in this figure we plot a histogram of 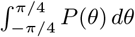 for all experiments except the two experiments for which winner/loser was declared inconclusive. Each experiment has two contributions to the histogram, one from the winner and one from the loser. Across all experiments, the 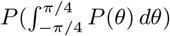 for winners and losers form two clearly distinct distributions. In each trial when assigning winner and loser, we are not simply choosing the larger of two similarly sized quantities. Rather, 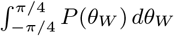 and 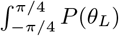 are robustly different in magnitude, showing that the symmetry between the contestants has been robustly broken, allowing for the labelling of winner and loser.

**FIG. S10.**
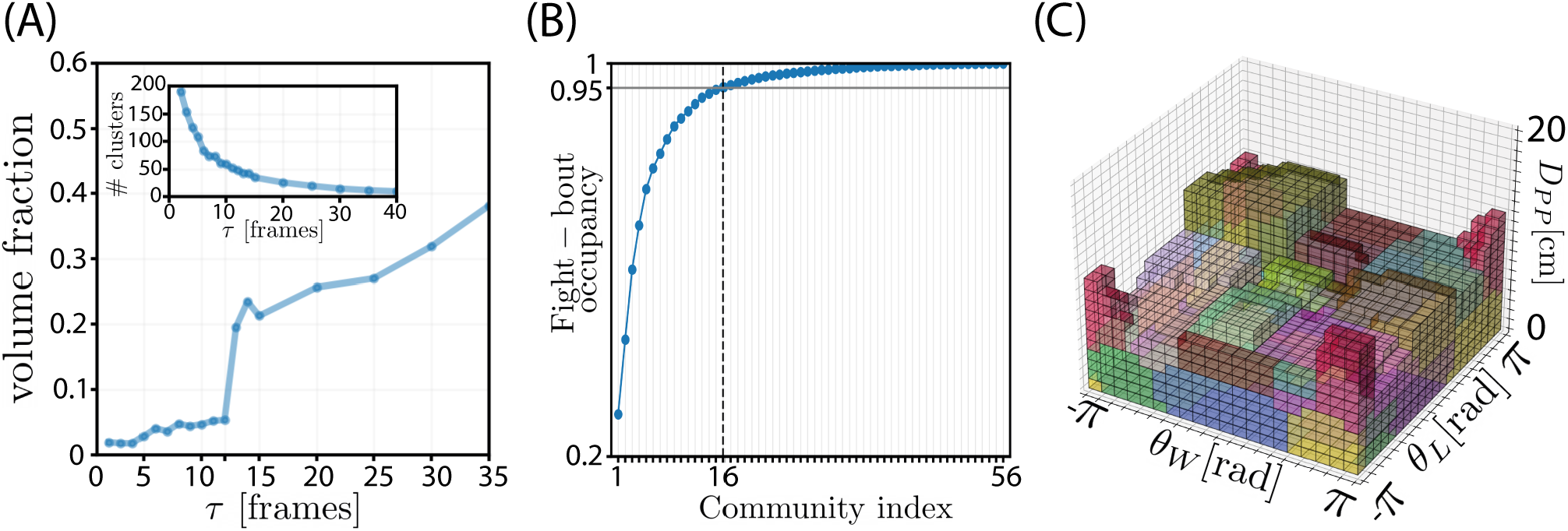
Determining the choice of *τ* for generating the (*D*_*PP*_, *θ*_*W*_, *θ*_*L*_) voxel transition matrix *T*_*ij*_, and identifying a subset of N=16 Infomap communities which span fight bouts. (A) We plot the volume fraction of the total voxel space spanned by the largest Infomap community as a function of *τ* . We see that as we approach *τ* = 12 frames, a transition appears where the largest community suddenly jumps in size. The Infomap communities which merge together are the close-distance (small *D*_*PP*_) communities, which are of particular interest. We interpret this as a loss of resolution of fight dynamics at *τ >* 12 frames. We hence chose *τ* below this transition, and set *τ* = 10 frames. (B, inset) We plot the number of Infomap communities as a function of *τ* . (B) When building *T*_*ij*_ we set the distance upper limit of *D*_*PP*_ = 20cm. Fighting generally occurs at *D*_*PP*_ ≪ 20cm, but we set this upper limit to avoid artificially truncating Infomap communities which have a large range in *D*_*PP*_ . Hence some of the N=56 Infomap communities identified from *T*_*ij*_ occur at larger distances than tend to occur during fighting. To focus on the maneuvers that occur during fight bouts, we sought to identify a subset of the N=56 Infomap communities which approximately span fight bouts. We sorted the the N=56 Infomap communities based on their cumulative probability in fight bouts, isolating a 16-member subset capturing 95% of the probability. (C) We show the full set of N=16 Infomap communities which are labelled as the fight maneuvers that span fight bouts (see Fig. 5 (A-middle)). We use N=16 colors meaning voxels which have the same coloring lie in the same Infomap community. The colors are drawn with some transparency to minimize the effect of occlusions among communities. The color-scheme for this panel does not match the color-scheme of Fig. 5 (A-right) and SI Fig. S11.

**FIG. S11.**
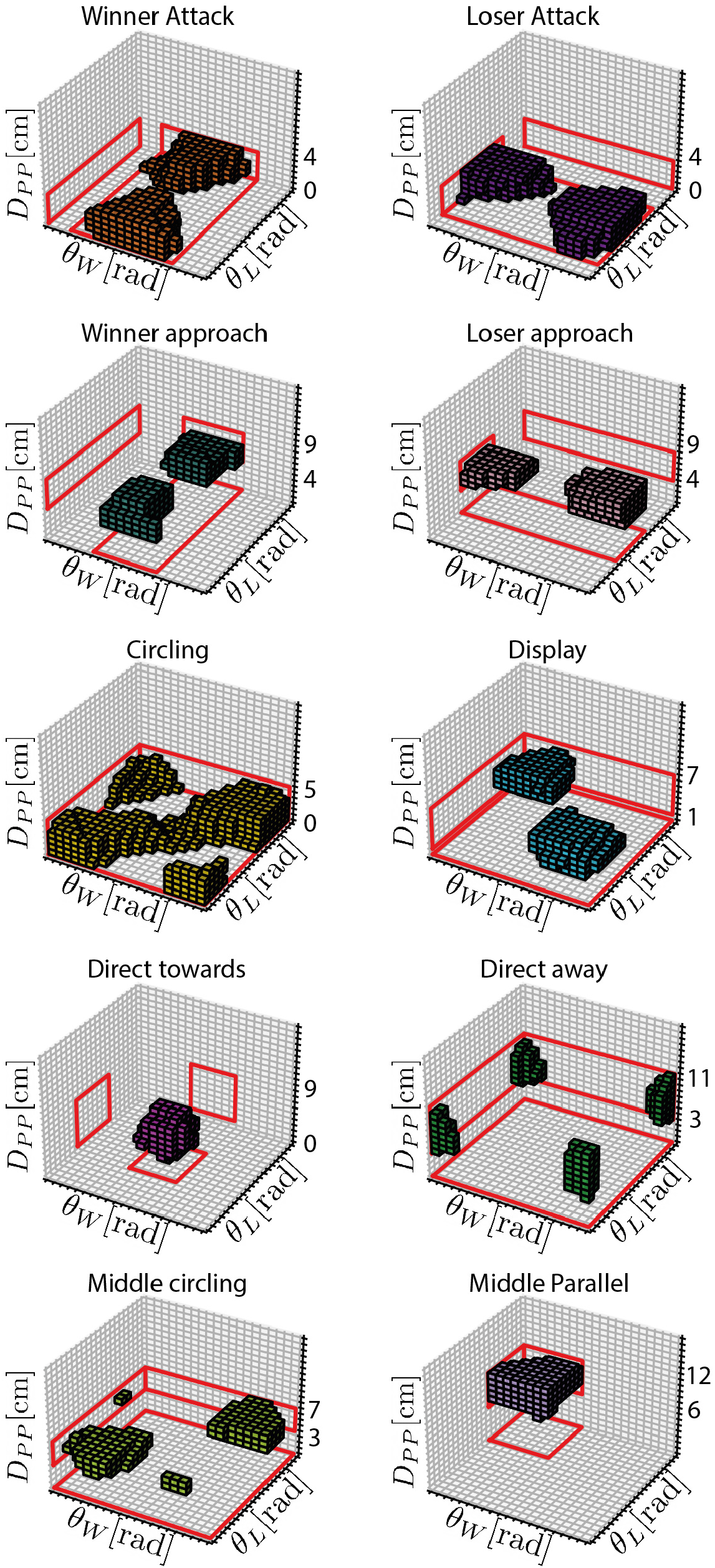
Expanded explanations of the fight maneuvers. Shown are the microstates which compose each of the N=10 fight maneuvers. The location of fight maneuvers in the configuration space allows us to interpret the maneuvers. See Methods for full explanation.

**FIG. S12.**
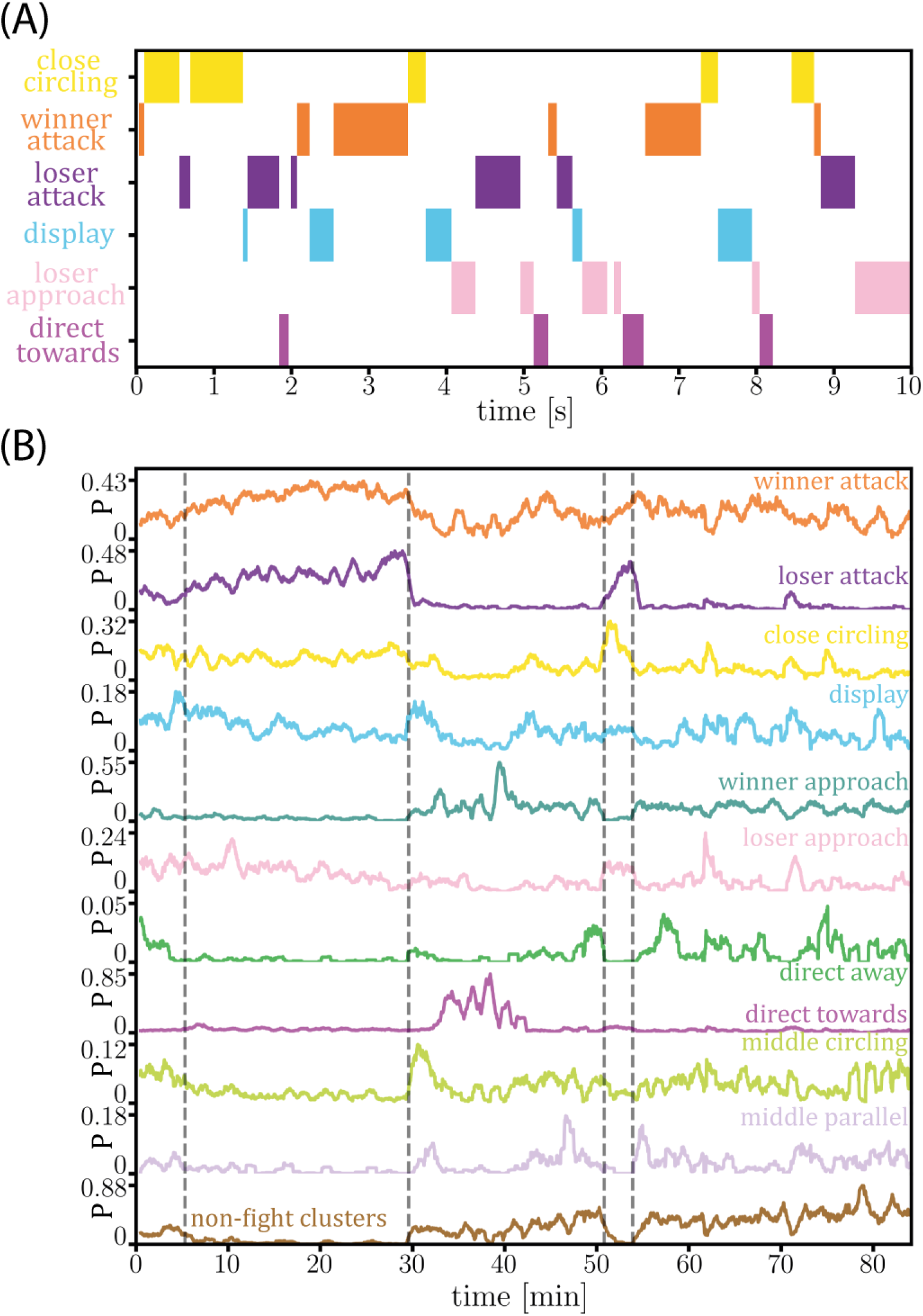
Changes in the probability of fight maneuvers reveal longer-time contest structure, including differential attack rates. (A) We show a sample ethogram composed of the fight maneuvers for a 10 s epoch during a fight bout. (B) Maneuver probabilities for a single imaging experiment. We show the 10 fight maneuvers, with the final row representing the probability of all non-fight maneuvers. Probabilities are estimated in 1-min running windows with 59-sec overlap. Grey lines denote temporal fight boundaries. During fight bouts, attack probabilities are high, and non-fight cluster probabilities are low. In this experiment, immediately after the first fight bout, the probability of maneuvers such as “display” and “direct towards” increases. These maneuvers suggest continued assessment and indeed there is a second fight bout later in the recording. At the beginning of the 2nd bout the loser attack probability rises sharply and when this bout concludes, the dominance decision has been reached; the pair visit asymmetric configurations favouring the winner almost all of the time for the remainder of the recording.

**FIG. S13.**
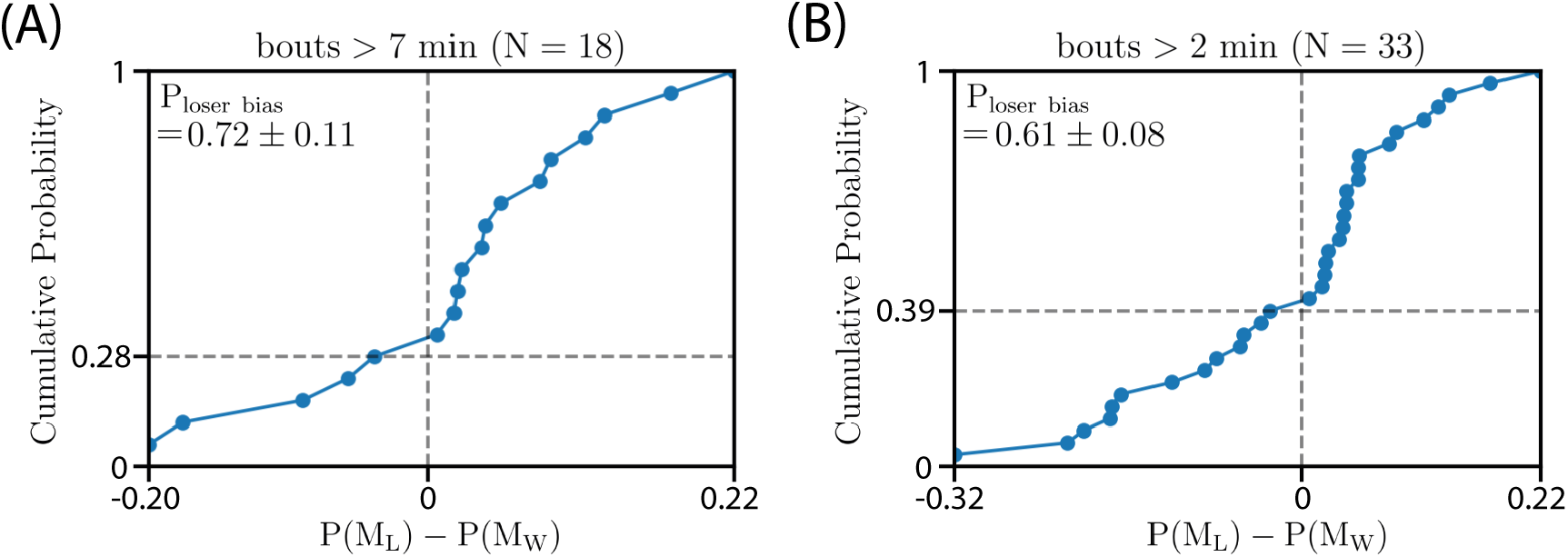
Across our full fight bout ensemble, there is a bias for bouts with more loser advantageous maneuvers than winner advantageous maneuvers in the last two minutes of the bout. We examine the statistics across individual bouts by computing the cumulative distribution of the difference in loser and winner advantageous maneuver probabilities in the last two minutes of each bout. (A) For the *N* = 18 bouts used in Fig. 6 (B) the probability of an individual bout having more loser advantageous maneuvers than winner advantageous maneuvers in the last two minutes is *P* _loser bias_ = 0.72 *±* 0.11, where the error is the standard deviation from bootstrapping across bouts. (B) This trend persists in our complete ensemble of *N* = 33 bouts (which are all longer than two minutes), *P* _loser bias_ = 0.61 0.08. We note that for shorter bouts the two-minute window over which we compute the maneuver probabilities is of a similar duration as the bout itself, blurring the difference between starting and ending behaviors, and thus expectedly reducing the loser advantageous maneuver bias.

SI Movie 1. **A visualization of the tracking pipeline output, namely the 3D bodypoint positions of two zebrafish over time, including organism identity**. In this animation, we show the 3D locations of the head (circle), pec (square) and tail (cross) bodypoints of the winner (red) and loser (blue) throughout an entire experiment (1hr 23min 29sec, FishTank20200130 153857), the same experiment shown in Fig. 3(A). The movie is downsampled from 100fps to 25fps to reduce file size.

SI Movie 2. **A visualization of the start of the first fight bout in FishTank20200130 153857**. In this movie we show two panels. The left panel shows an animation of the contestant 3D bodypoint positions. The right panel shows the XZ camera view of the tank with the XZ image coordinates of bodypoints drawn over the fish. The movie is 2 minutes long, and runs from one minute before until one minute after the frame identified as the start of the first fight bout.

SI Movie 3. **A visualization of the middle of the first fight bout in FishTank20200130 153857**. In this movie we show two panels. The left panel shows an animation of the contestant 3D bodypoint positions. The right panel shows the XZ camera view of the tank with the XZ image coordinates of bodypoints drawn over the fish. The movie is 2 minutes long, and runs from one minute before until one minute after the frame identified as the middle of the first fight bout.

SI Movie 4. **A visualization of the end of the first fight bout in FishTank20200130 153857**. In this movie we show two panels. The left panel shows an animation of the contestant 3D bodypoint positions. The right panel shows the XZ camera view of the tank with the XZ image coordinates of bodypoints drawn over the fish. The movie is 2 minutes long, and runs from one minute before until one minute after the frame identified as the end of the first fight bout.

SI Movie 5. **A visualization of the start of the second fight bout in FishTank20200130 153857**. In this movie we show two panels. The left panel shows an animation of the contestant 3D bodypoint positions. The right panel shows the XZ camera view of the tank with the XZ image coordinates of bodypoints drawn over the fish. The movie is 2 minutes long, and runs from one minute before until one minute after the frame identified as the start of the second fight bout.

SI Movie 6. **A visualization of the end of the second fight bout in FishTank20200130 153857**. In this movie we show two panels. The left panel shows an animation of the contestant 3D bodypoint positions. The right panel shows the XZ camera view of the tank with the XZ image coordinates of bodypoints drawn over the fish. The movie is 2 minutes long, and runs from one minute before until one minute after the frame identified as the end of the second fight bout.

